## Supplemental figures for "Cell type-specific *cis*-regulatory divergence in gene expression and chromatin accessibility revealed by human-chimpanzee hybrid cells"

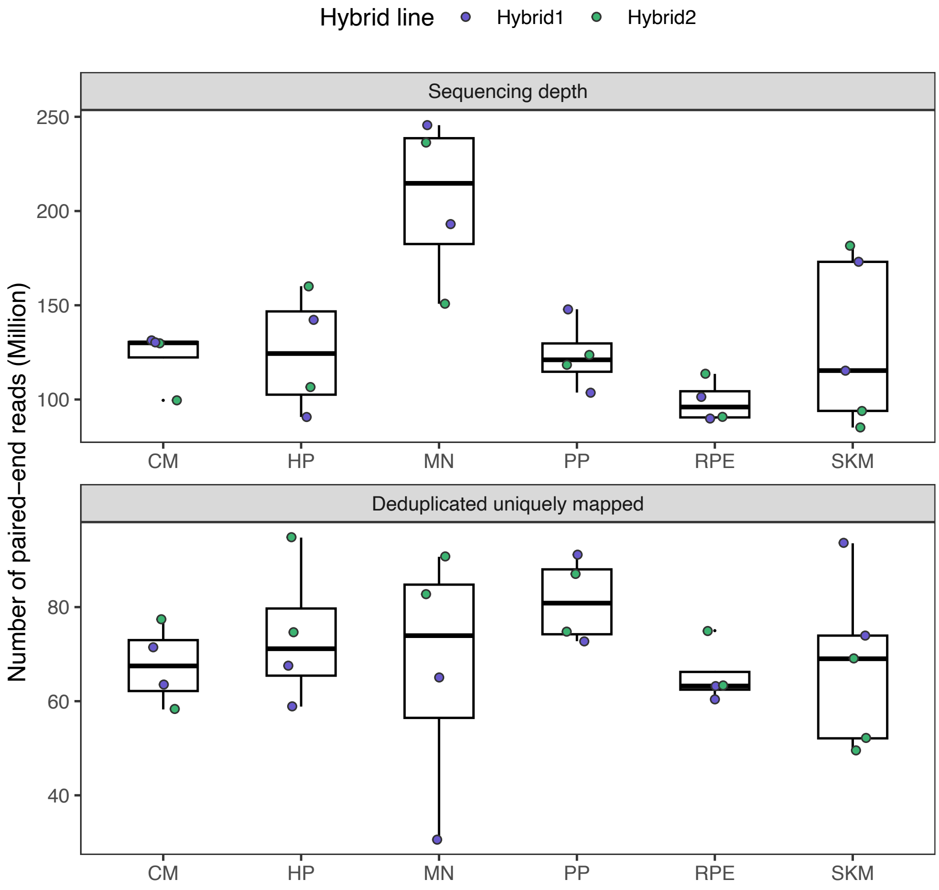


**Supplemental Figure 1. Sequencing depth across samples for the RNA-seq data.** Sequencing depth in the raw fastq files is shown in the top panel. The bottom panel shows the read count after filtering for only uniquely mapped reads and removing duplicates. CM: cardiomyocyte; HP: hepatocyte progenitor; MN: motor neuron; PP: pancreatic progenitor; RPE: retinal pigment epithelium; SKM: skeletal myocyte.

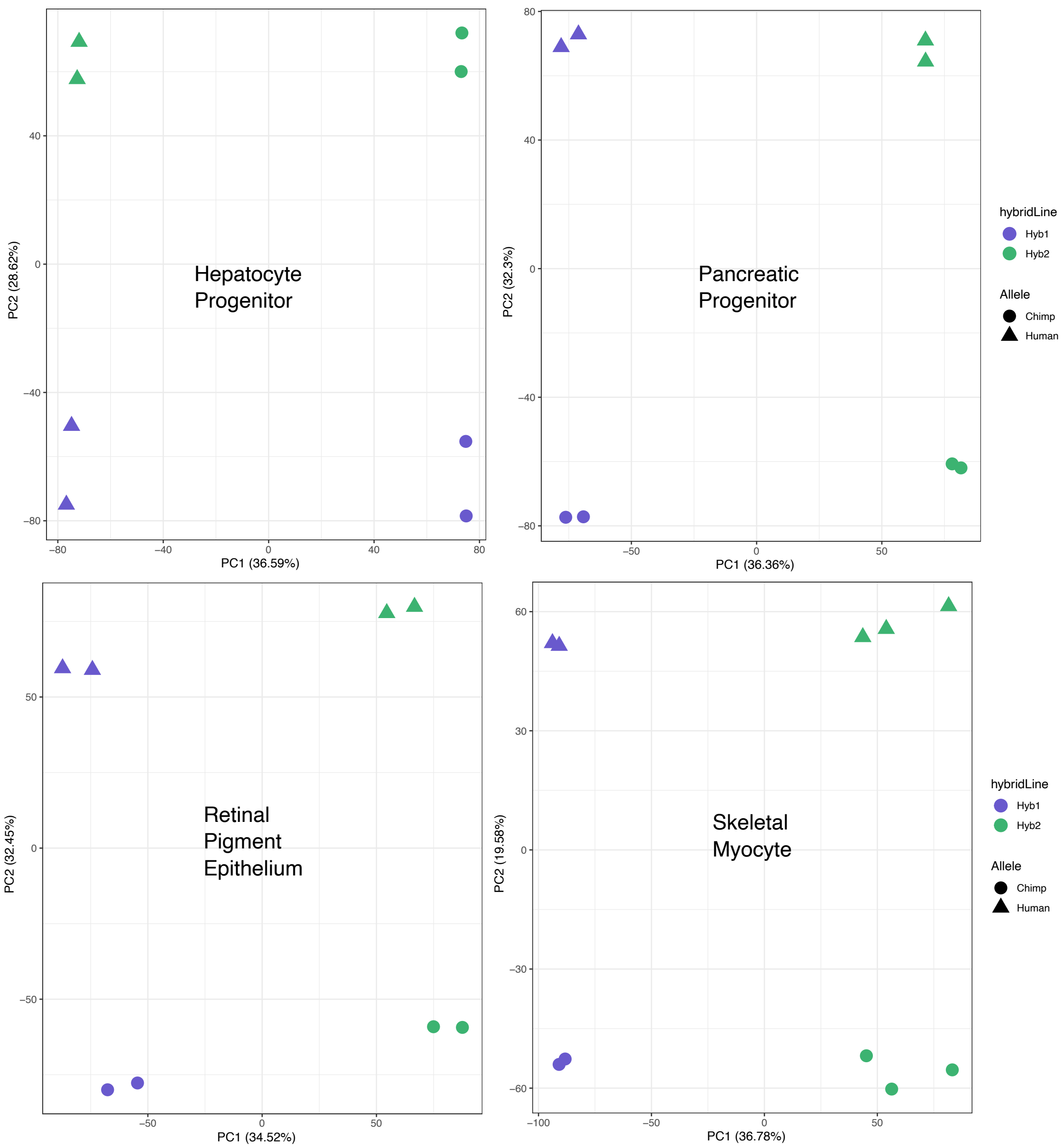


**Supplemental Figure 2. PCA on allelic counts from RNA-seq for individual cell types.** The result of running principal component analysis (PCA) on the individual cell types not shown in Fig. 1d. Species were clearly separated by PC1 or PC2 when PCA was performed on each cell type.

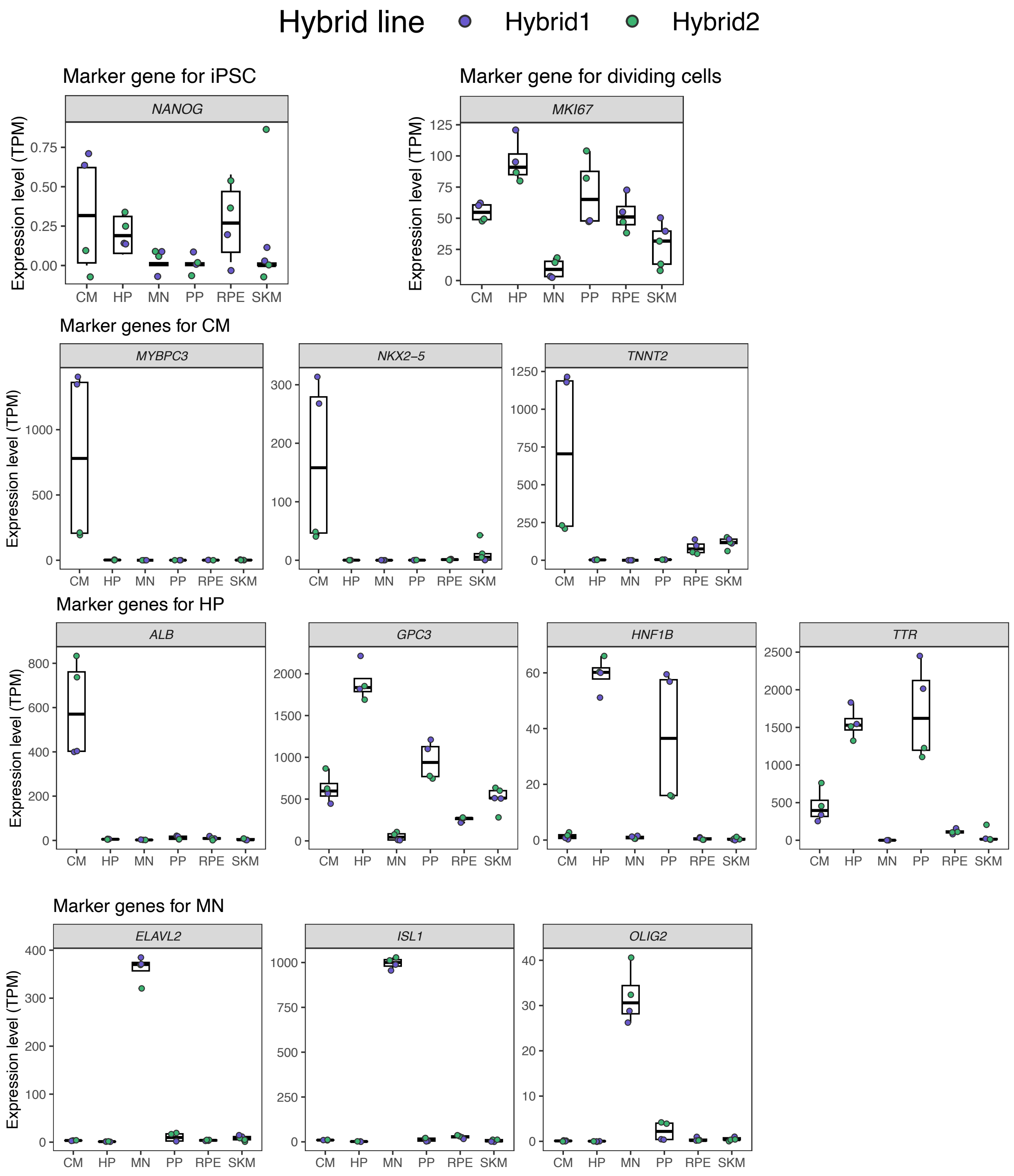
Hybrid line
Hybrid1
Hybrid2
Marker gene for iPSC
NANOG
Marker gene for dividing cells
MKI67
Marker genes for CM
MYBPC3
NKX2–5
TNNT2
Marker genes for HP
ALB
GPC3
HNF1B
TTR
Marker genes for MN
ELAVL2
ISL1
OLIG2
Expression level (TPM)
CM HP MN PP RPE SKM

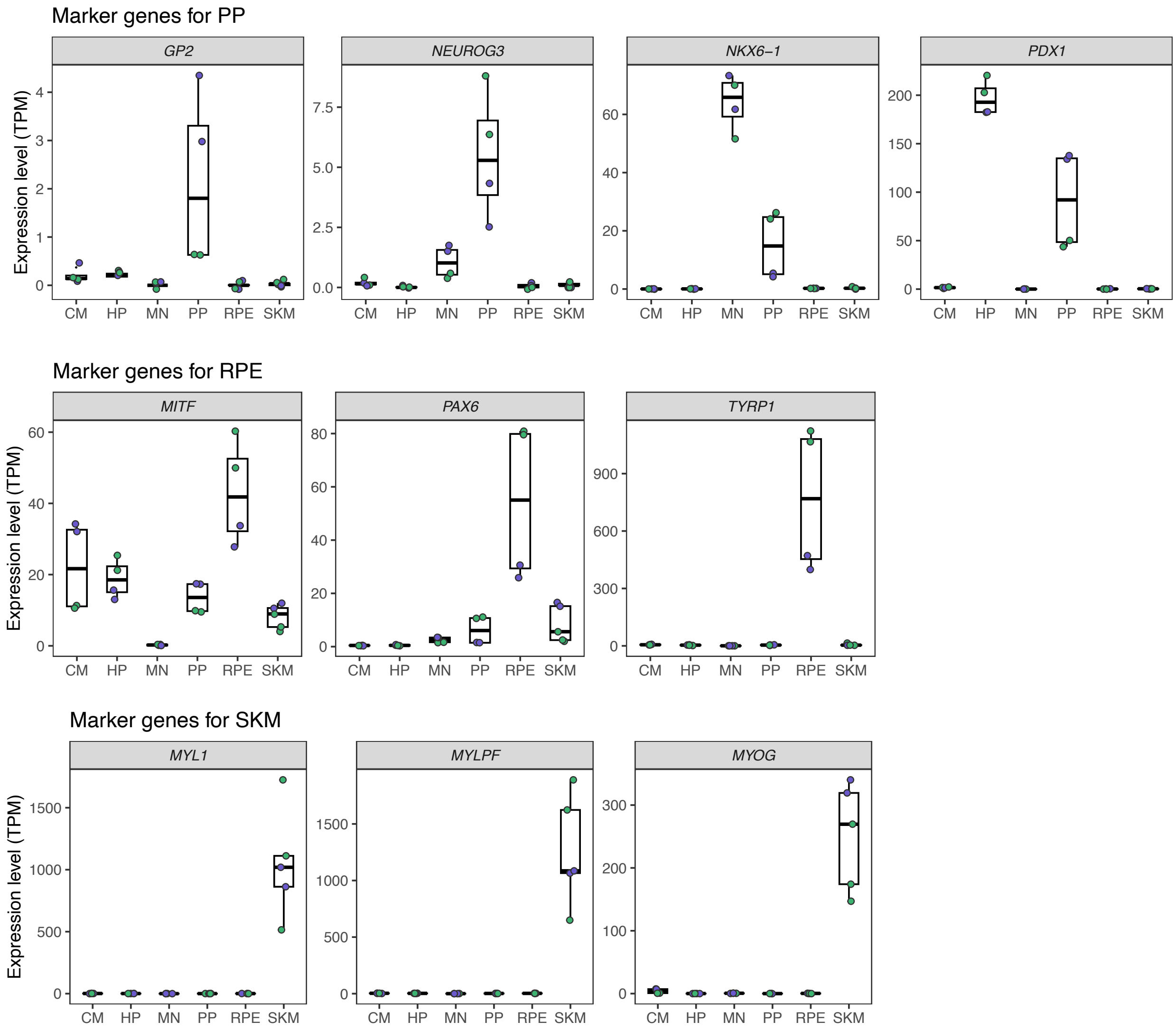


**Supplemental Figure 3. Marker gene expression in different cell types.** In order, the panels show: a marker for pluripotency, a marker gene for dividing cells, marker genes for cardiomyocytes, marker genes for hepatocytes and hepatocyte progenitors, marker genes for motor neurons, marker genes for pancreatic progenitors and more mature pancreatic cell types, marker genes for retinal pigment epithelial cells, and marker genes for skeletal myocytes. Hepatocyte progenitors and pancreatic progenitors generally show similar gene expression profiles. TPM: transcript per million.

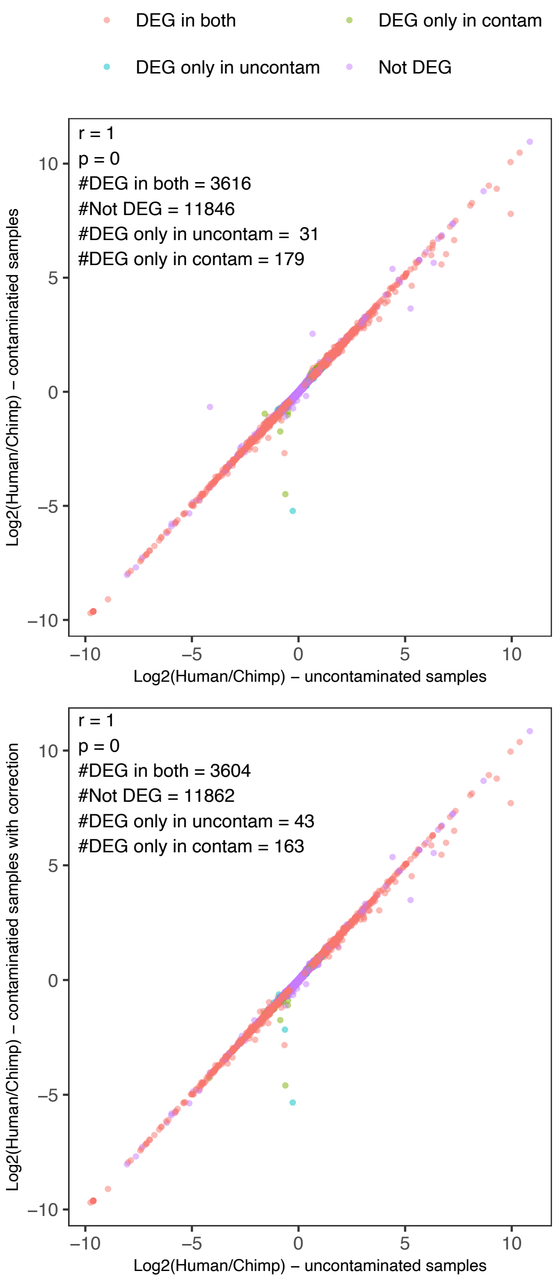


**Supplemental Figure 4. Simulated chimpanzee parental contamination of hybrid RNA-seq data and correction.** The result of using hybrid and chimpanzee parental iPSC RNA-seq from Agoglia et al. (the same lines used in this study) is shown. Simulation of chimpanzee parental contamination was done by mixing chimpanzee parental data into hybrid2 data to reach a similar chimpanzee bias to that observed in PP hybrid2 samples. Top: $\log_2$(human allele counts/chimpanzee allele counts) estimates from uncontaminated samples vs contaminated samples. Bottom: $\log_2$(human allele counts/chimpanzee allele counts) estimates from uncontaminated samples vs contaminated samples after correction. The numbers of detected differentially expressed genes with FDR<0.05 in each condition are shown on the figures.

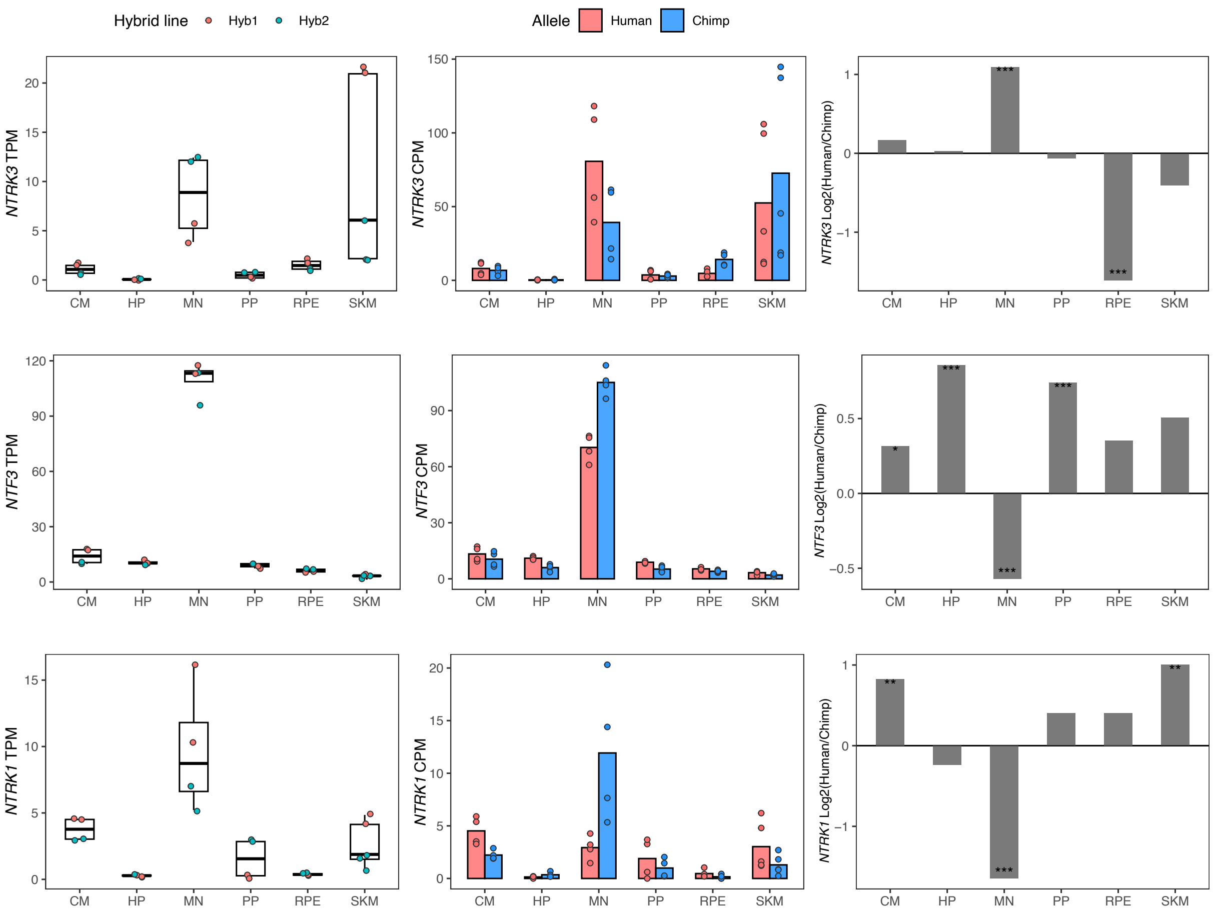


**Supplemental Figure 5. *NTF3*, *NTRK1* and *NTRK3* have cell type-specific allele-specific expression.** Left: TPM from total expression, with the mean TPM as the center of boxplots and the TPM of each sample plotted as dots. Samples were colored by the hybrid line from which they were collected. Middle: Allelic counts per million (CPM). Right: DESeq2 log fold-changes ($\log_2$(human/chimpanzee)) and significance (using DESeq2 likelihood ratio test) which is indicated by asterisks where *** indicates FDR < 0.005, ** indicates FDR < 0.01, and * indicates FDR < 0.05.

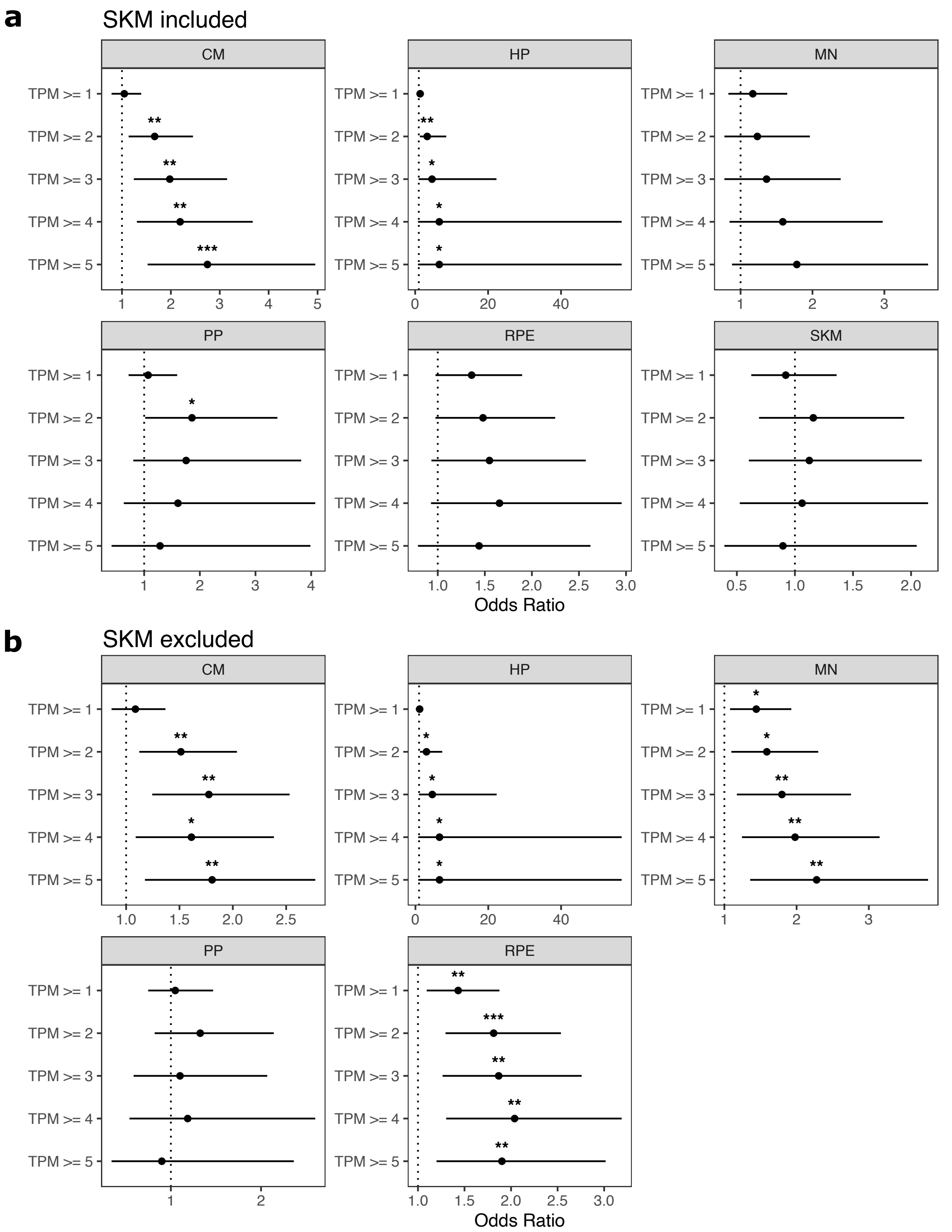


**Supplemental Figure 6. Genes expressed in only one cell type are enriched for genes with ASE. a)** TPM cutoffs were used to test whether genes expressed in only one cell type are enriched for genes with allele-specific expression (ASE). For each tested cell type, genes with TPM greater than the cutoff in this cell type and TPM < 1 in all other cell types were tested. Significance (using the Wald test) is indicated by asterisks where *** indicates $p < 0.005$, ** indicates $p < 0.01$, and * indicates $p < 0.05$. **b)** The same as in (a) but excluding SKM.

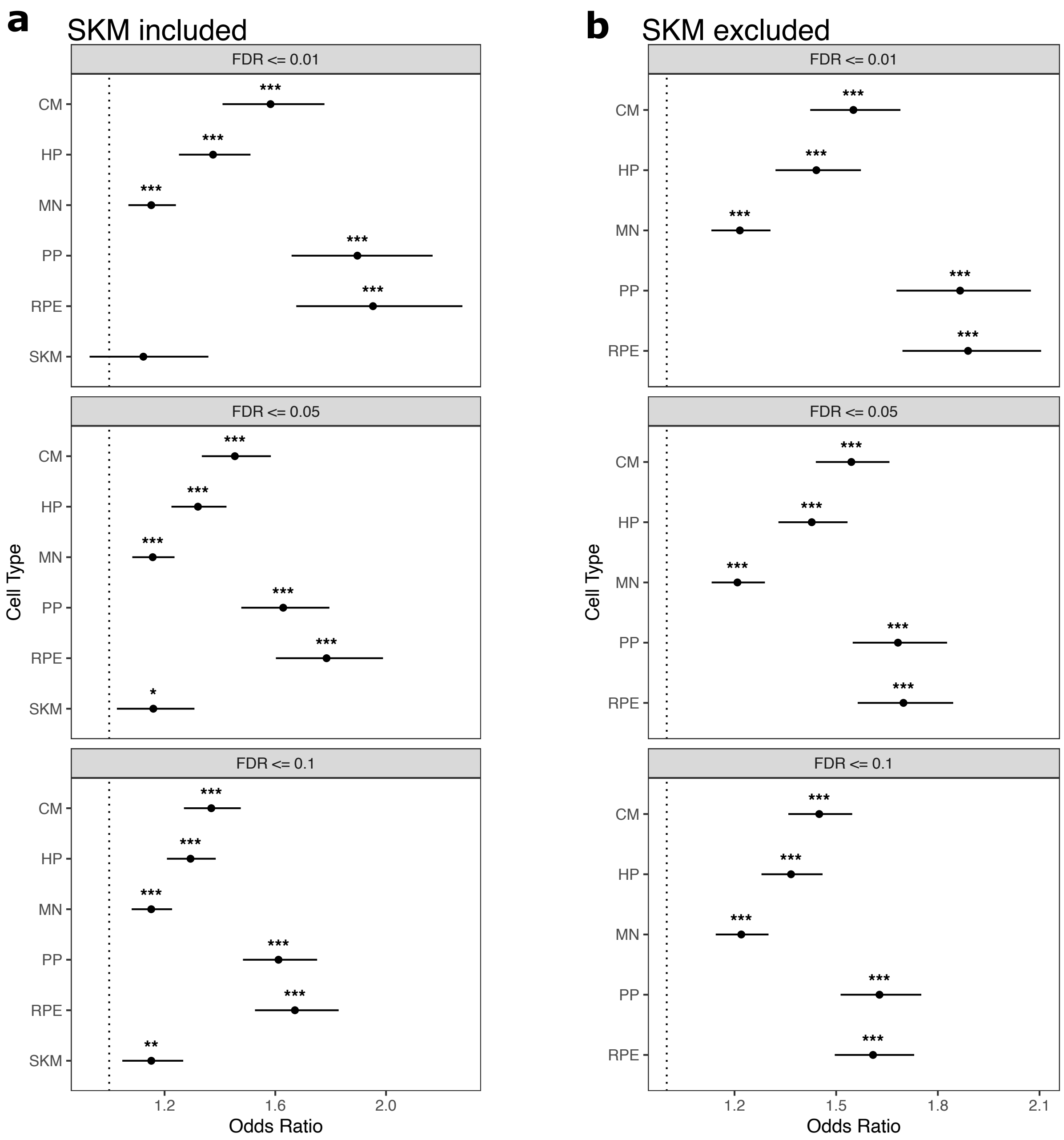


**Supplemental Figure 7. Cell type-specifically expressed genes are enriched for genes with ASE across FDR cutoffs. a)** Testing for enrichment of tissue-specific genes for ASE genes across various FDR cutoffs. Significance (using the Wald test) is indicated by asterisks where *** indicates $p < 0.005$, ** indicates $p < 0.01$, and * indicates $p < 0.05$. **b)** The same as in (a) but excluding SKM.

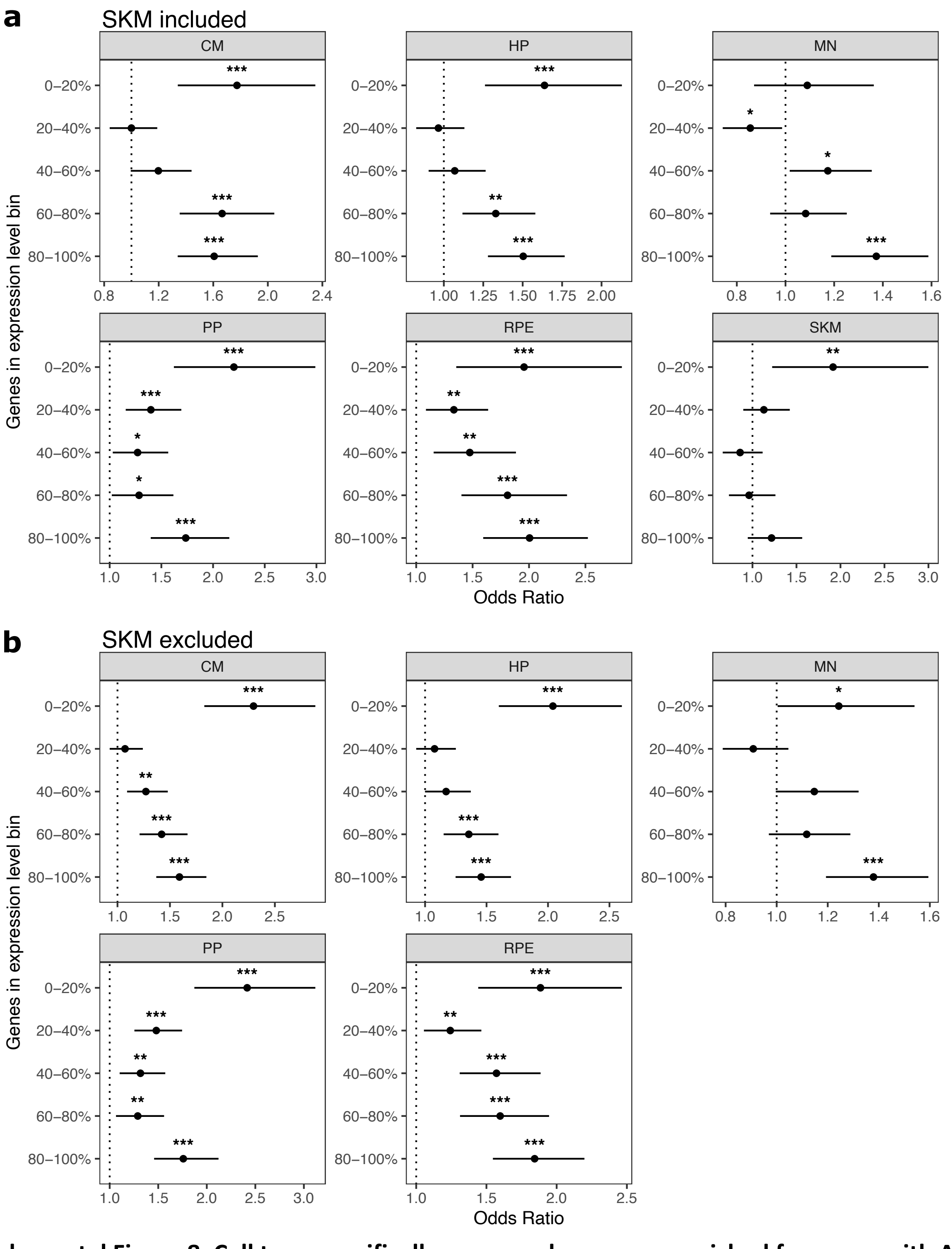


**Supplemental Figure 8. Cell type-specifically expressed genes are enriched for genes with ASE regardless of expression level. a)** The enrichment for cell type-specific genes in genes with ASE holds regardless of gene expression level. Significance (using the Wald test) is indicated by asterisks where *** indicates $p < 0.005$, ** indicates $p < 0.01$, and * indicates $p < 0.05$. Genes in the 0-20% bin are the most lowly expressed whereas genes in the 80-100% bin are the most highly expressed. **b)** The same as in (a) but excluding SKM.

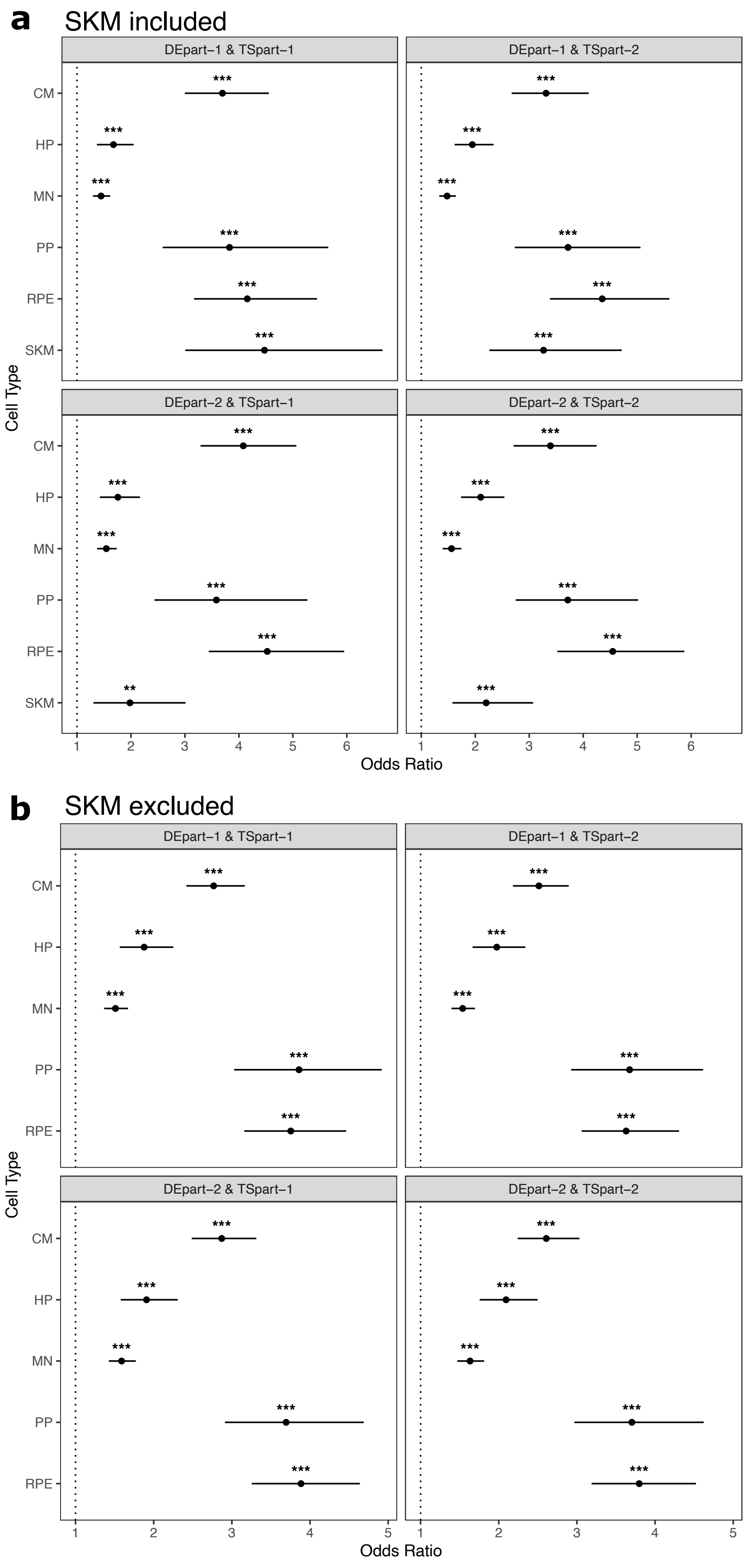
a
SKM included
DEpart−1 & TSpart−1
DEpart−1 & TSpart−2
DEpart−2 & TSpart−1
DEpart−2 & TSpart−2
CM
HP
MN
PP
RPE
SKM
***
**
Cell Type
Odds Ratio
b
SKM excluded
DEpart−1 & TSpart−1
DEpart−1 & TSpart−2
DEpart−2 & TSpart−1
DEpart−2 & TSpart−2
CM
HP
MN
PP
RPE
Cell Type
Odds Ratio

**Supplemental Figure 9. Cell type-specifically expressed genes are enriched for genes with ASE when different samples are used to identify cell type-specific and ASE genes. a)** At least four samples including 2 hybrid1 samples and 2 hybrid2 samples were collected for each cell type, except 5 samples were collected for SKM (2 hybrid1 and 3 hybrid2 samples). Samples were split into part-1 and part-2 randomly by enforcing that each part have both hybrid1 and hybrid2 samples. Genes with ASE (denoted here as DE) and tissue-specifically expressed genes (denoted here as TS) were identified from part-1 and part-2 respectively (or vice versa), so different samples were used for each definition. For example, DEpart-1 indicates that genes with ASE were identified from part-1 samples and TSpart-2 indicates that tissue- specifically expressed genes were identified from part-2 samples. Significance (using the Wald test) is indicated by asterisks where *** indicates $p < 0.005$, ** indicates $p < 0.01$, and * indicates $p < 0.05$. **b)** The same as in (a) but excluding SKM.

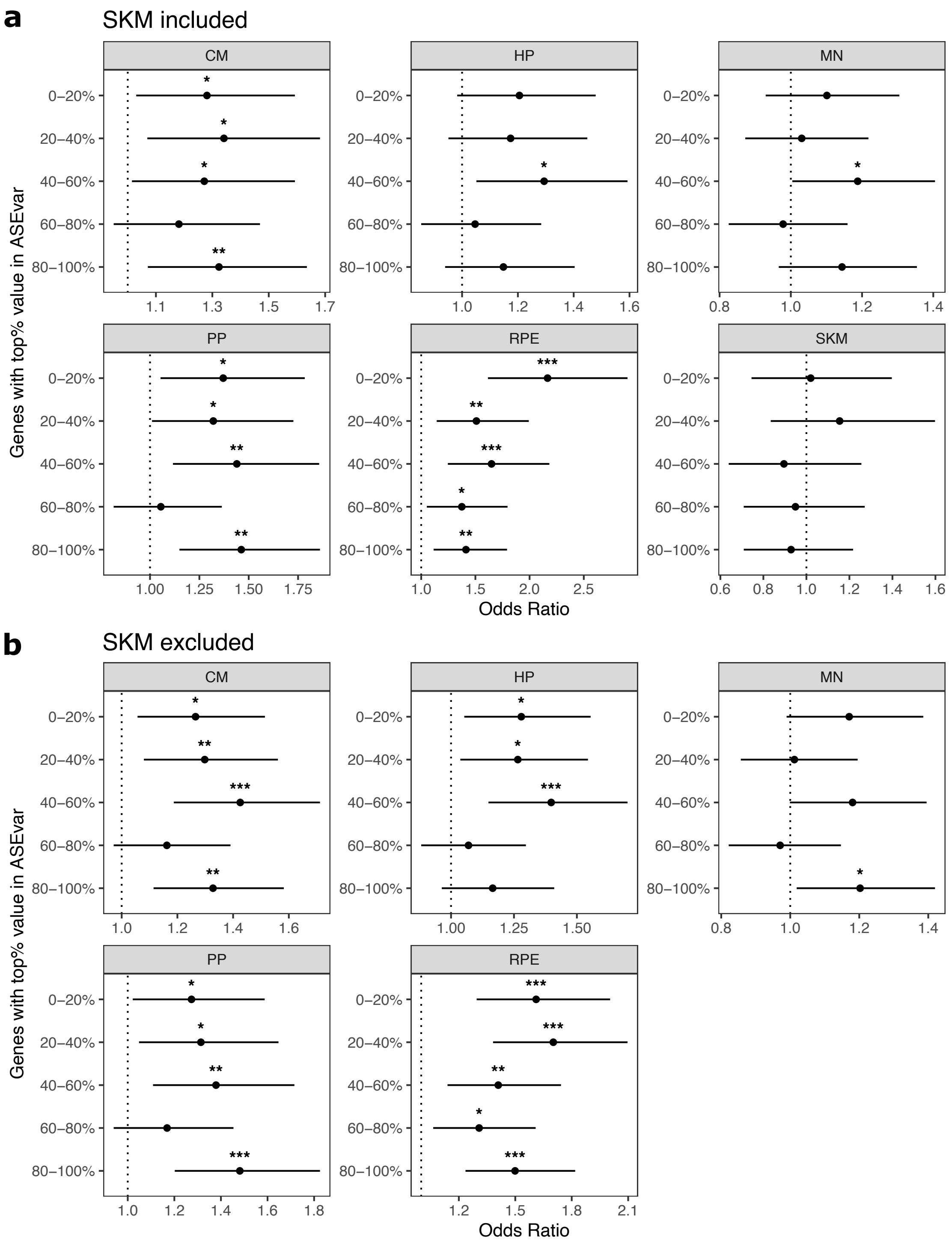


**Supplemental Figure 10. Controlling for ASE variance generally does not eliminate enrichment of ASE genes in cell type-specifically expressed genes. a)** Enrichments tests when controlling for ASE variance, a metric that measures constraint on gene expression developed by Starr et al. Genes were split into equal size bins based on ASE variance values. Genes in the 0-20% bin are the most constrained while genes in the 80-100% bin are the least constrained. Significance (using the Wald test) is indicated by asterisks where *** indicates $p < 0.005$, ** indicates $p < 0.01$, and * indicates $p < 0.05$. **b)** The same as in (a) but excluding SKM.

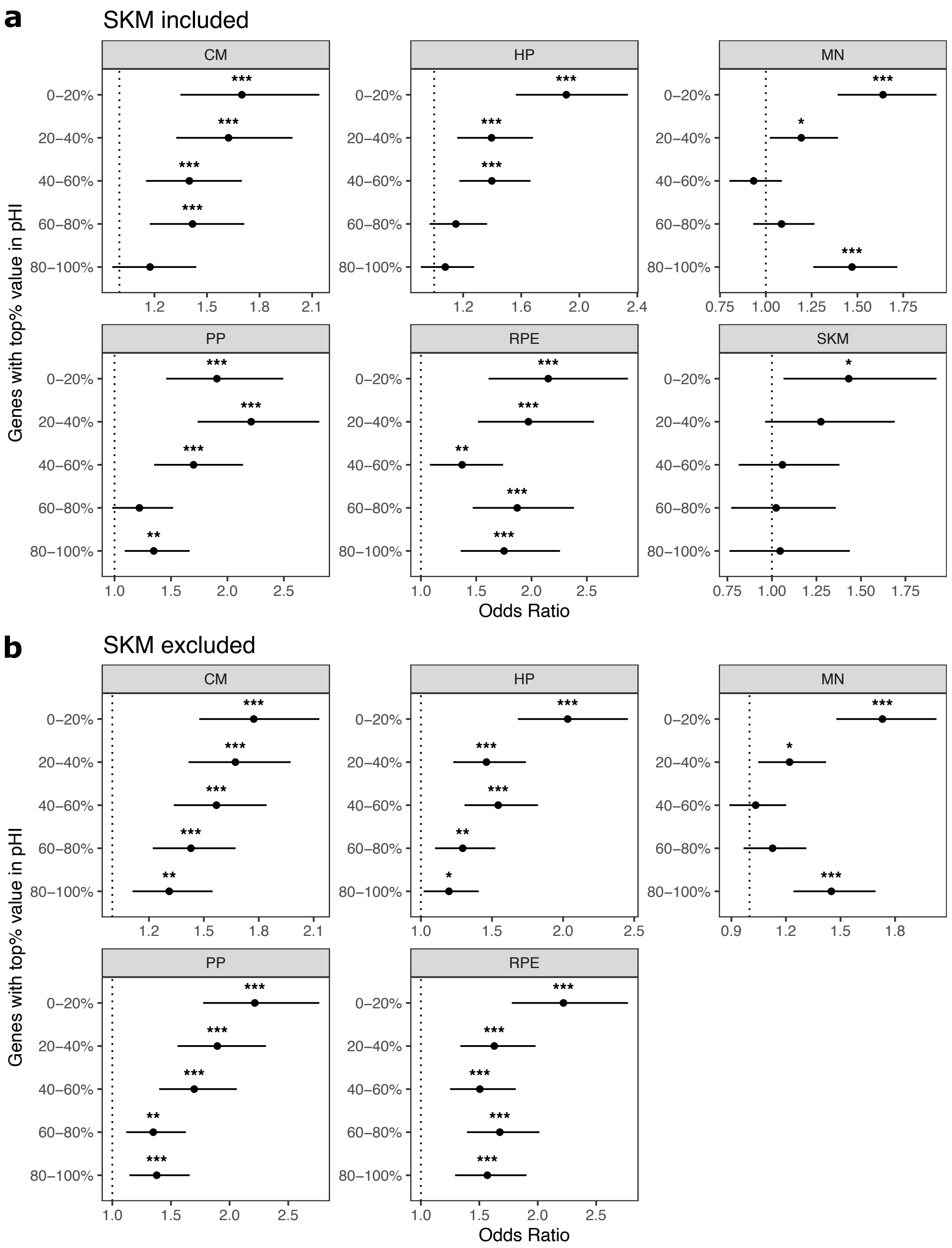


**Supplemental Figure 11. Controlling for probability of haploinsufficiency (pHI) generally does not eliminate enrichment of ASE genes in cell type-specifically expressed genes. a)** Enrichment tests when controlling for the probability of haploinsufficiency score (pHI) score. Genes were split into equal size bins based on the pHI values. Genes in the 80-100% bin are the most constrained while genes in the 0-20% bin are the least constrained. Significance (using the Wald test) is indicated by asterisks where *** indicates $p < 0.005$, ** indicates $p < 0.01$, and * indicates $p < 0.05$. **b)** The same as in (a) but excluding SKM.

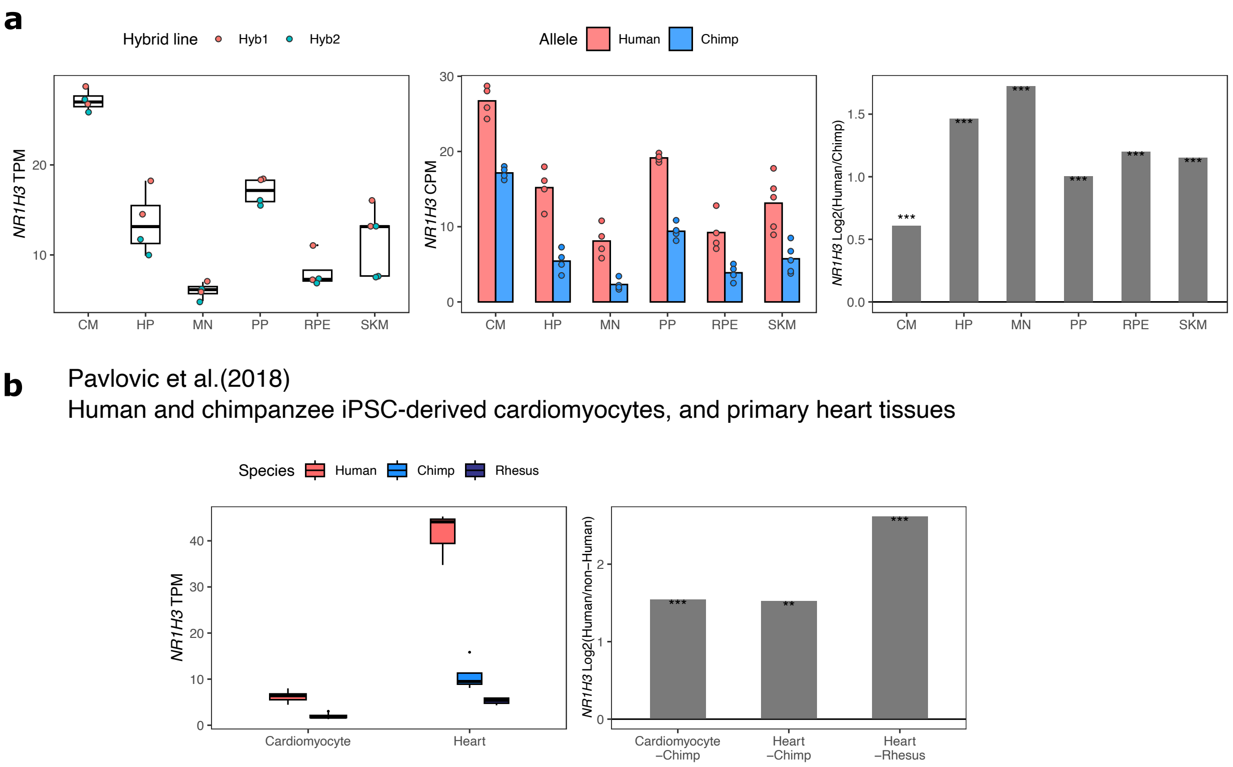


**Supplemental Figure 12. Validation of increased *NR1H3* expression in the human lineage. a)** TPM, allelic CPM, and $\log_2$ fold-change estimated by DESeq2 across cell types in our dataset for *NR1H3*. Significance (using the DESeq2 likelihood ratio test) is indicated by asterisks where *** indicates FDR < 0.005, ** indicates FDR < 0.01, and * indicates FDR < 0.05. **b)** *NR1H3* expression and estimated log fold-change in human and chimpanzee cardiomyocytes and human, chimpanzee, and rhesus macaque adult heart tissue. These results indicate that the increase in *NR1H3* expression occurred on the human lineage.

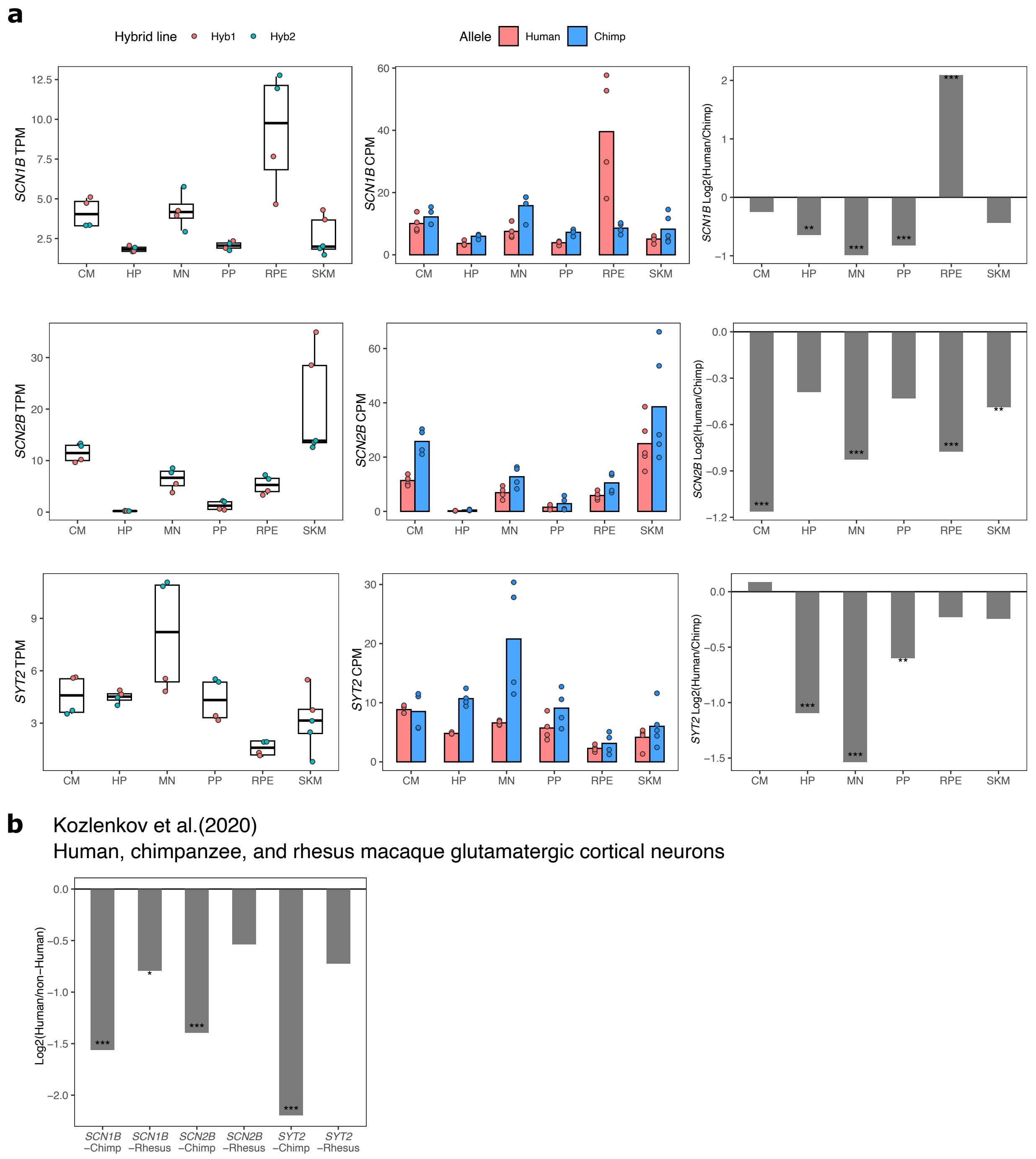


**Supplemental Figure 13. ASE and differential expression of *SCN1B*, *SCN2B*, and *SYT2* across cell types. a)** TPM, allelic CPM, and $\log_2$ fold change estimated by DESeq2 for *SCN1B*, *SCN2B*, and *SYT2* across cell types in our dataset. Significance (using the DESeq2 likelihood ratio test) is indicated by asterisks where *** indicates FDR < 0.005, ** indicates FDR < 0.01, and * indicates FDR < 0.05. **b)** $\mathrm{Log}_2$ fold-changes and FDR for *SCN1B*, *SCN2B*, *SYT2* in cortical glutamatergic neurons as computed by Kozlenkov et al.

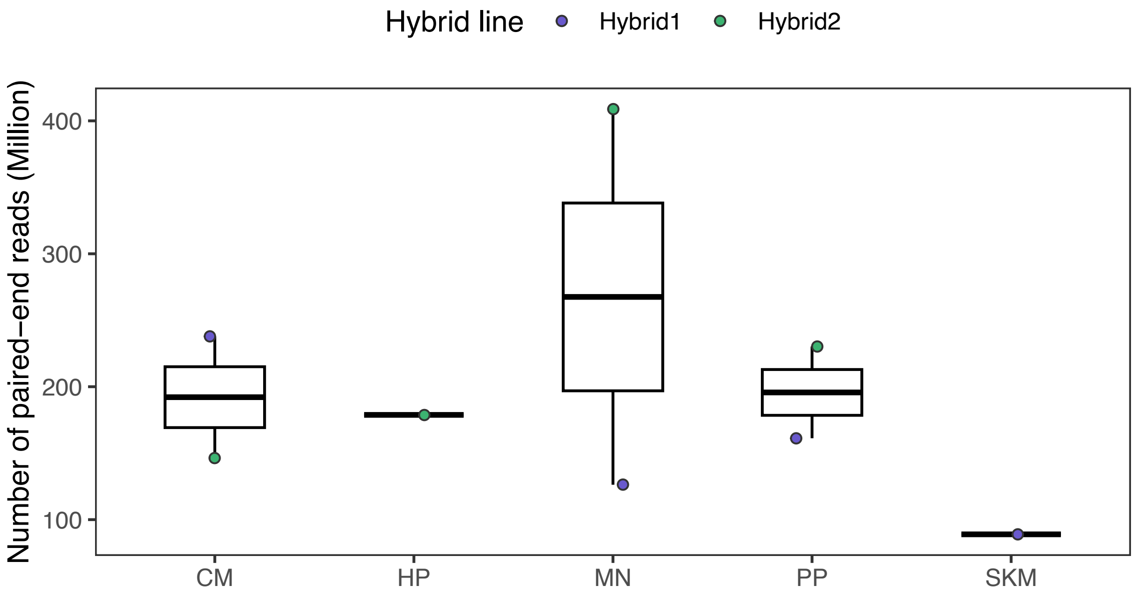


**Supplemental Figure 14. Sequencing depth in the ATAC-seq dataset.** Raw read counts from fastq files are shown. CM, MN, and PP have two replicates. ATAC: assay for transposase accessible chromatin.

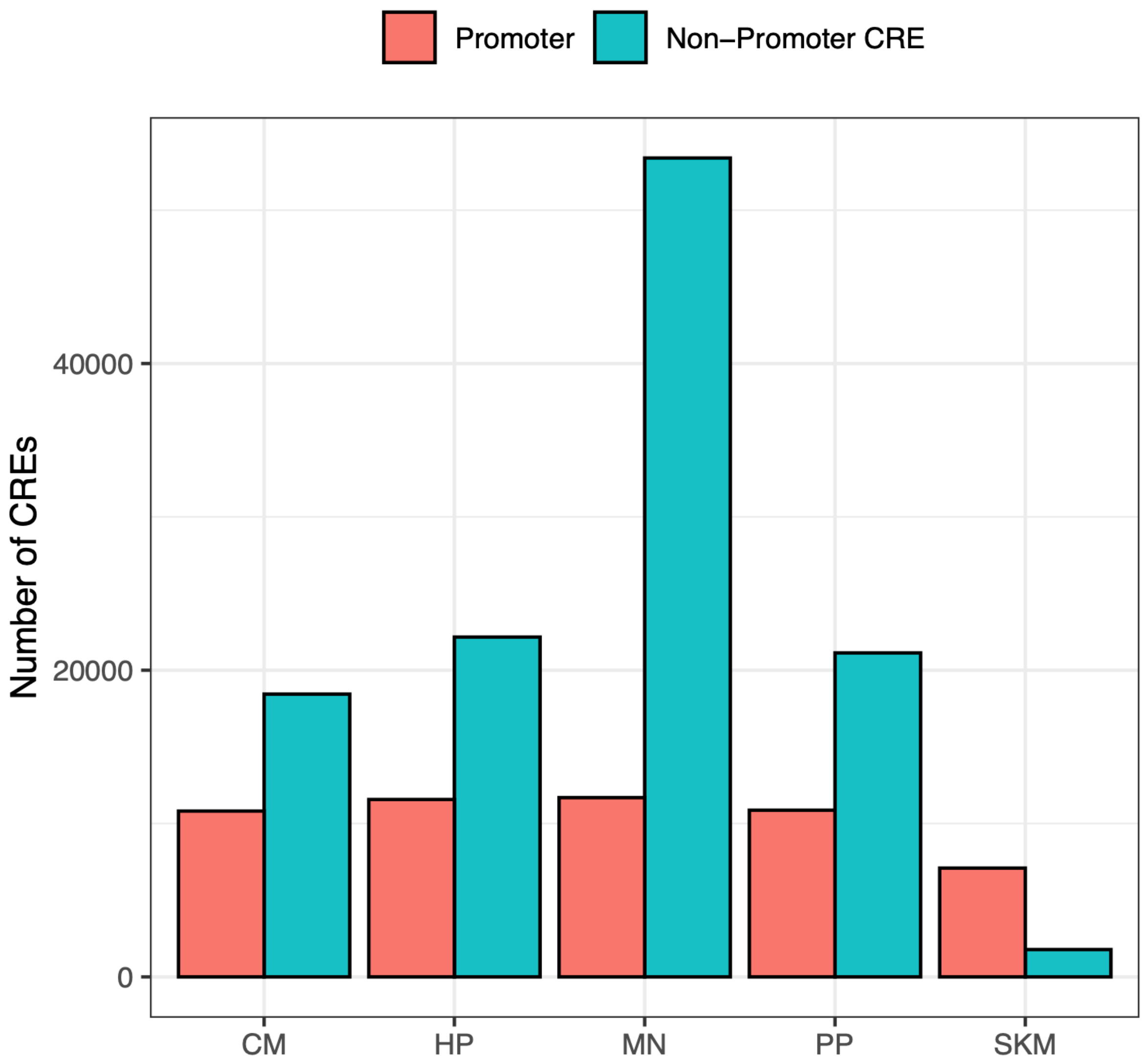


**Supplemental Figure 15**. **Number of CREs identified in each cell type.** ATAC-seq peaks from each cell type were categorized as 'Promoter' and 'Non-Promoter CRE' based on whether peaks overlapped with a TSS (see Methods). The high number in MN and low number in SKM are likely due to sequencing depth differences.

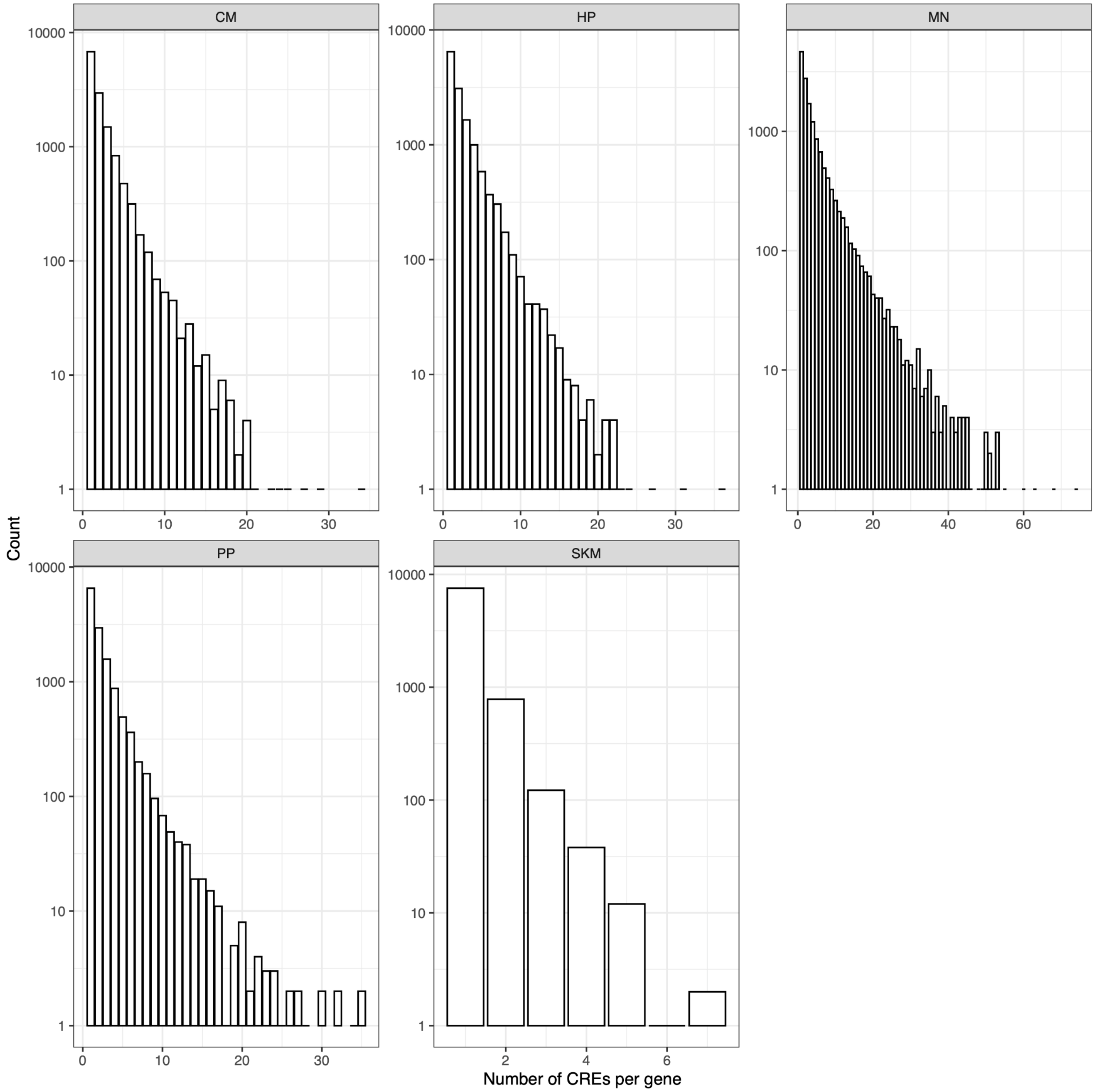


**Supplemental Figure 16**. **Number of CREs per gene identified in each cell type.** ATAC-seq peaks were assigned to the closest gene. The histogram of the number of CREs per gene is shown and note the y axis is on log scale.

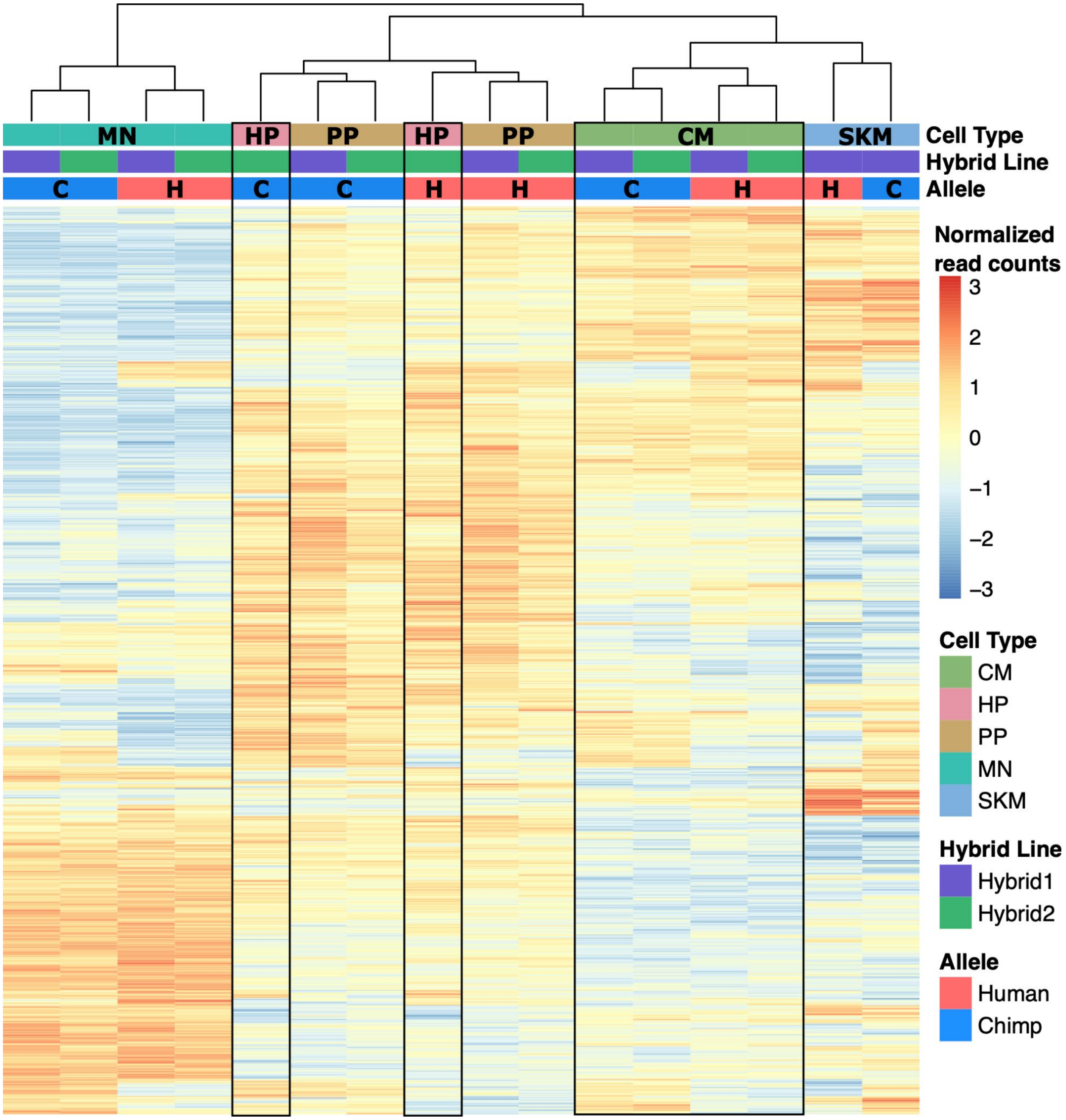


**Supplemental Figure 17. Heatmap and clustering of ATAC-seq samples.** Heatmap and result of hierarchical clustering performed on normalized allelic counts for the ATAC-seq dataset. Samples primarily clustered by cell type except HP and PP which clustered by allele. H: human; C: chimp.

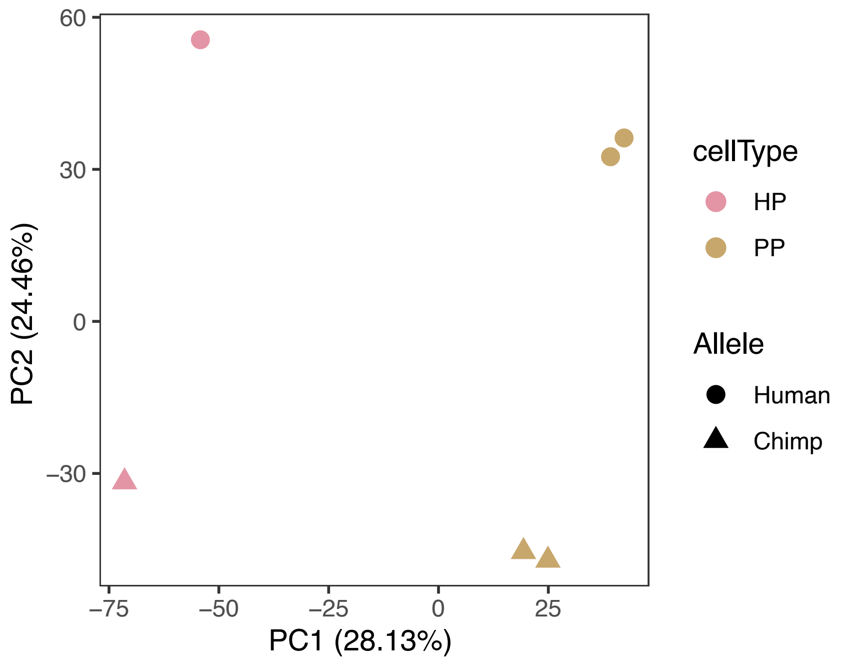


**Supplemental Figure 18. PCA of HP and PP ATAC-seq samples only**. Cell types are clearly separated by PC1 and species were clearly separated by PC2.

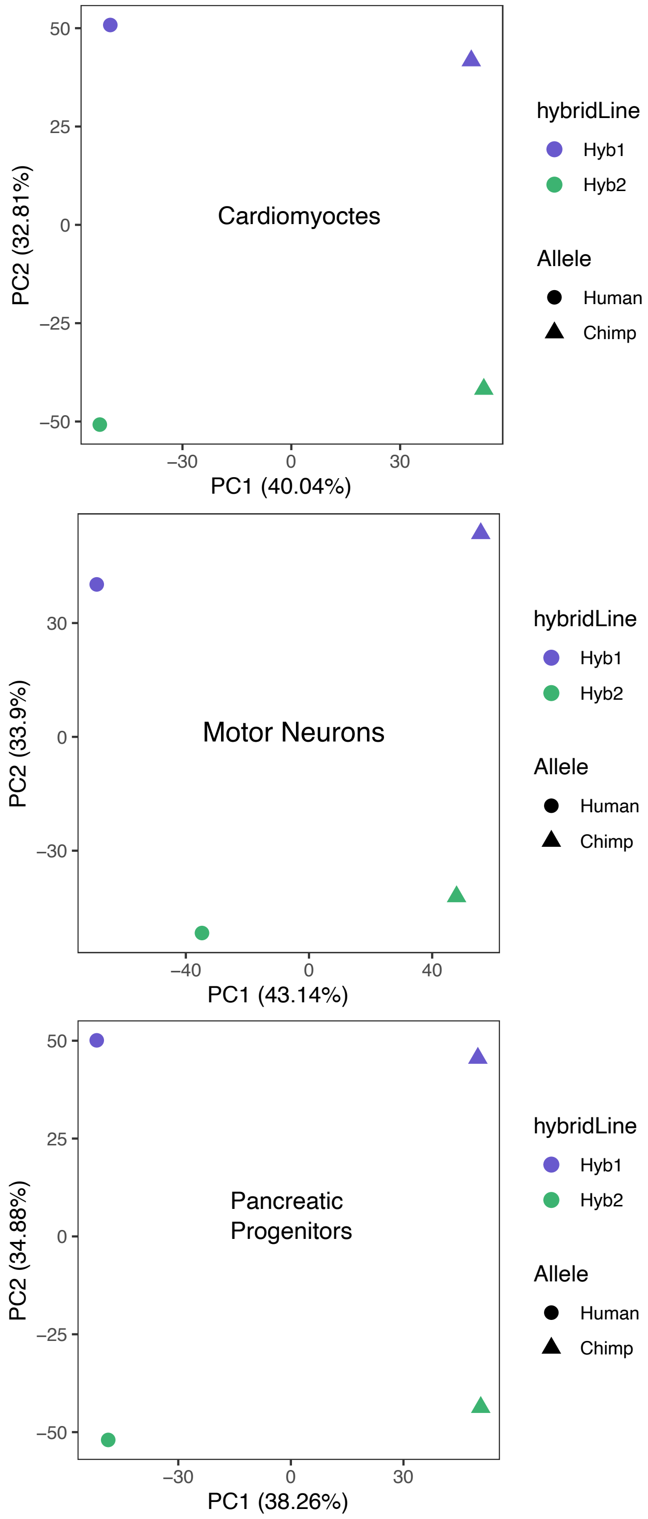


**Supplemental Figure 19. PCA separates the human and chimpanzee alleles in the ATAC-seq data.** PCA on allelic counts from ATAC-seq for each cell type that had more than one replicate is shown. Species are clearly separated by PC1 or PC2 when PCA was performed on each cell type.

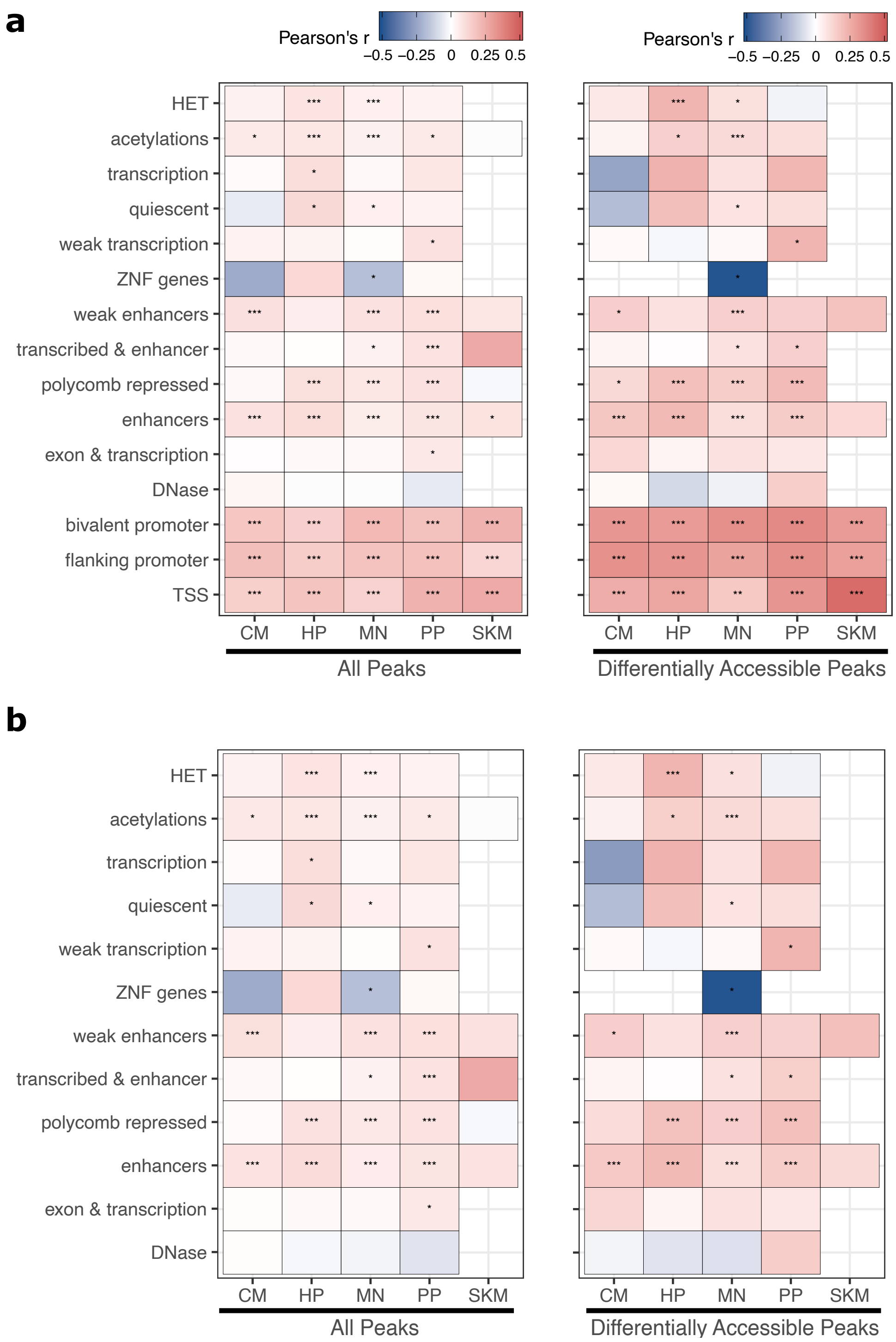


**Supplemental Figure 20. ASCA predicts ASE across a variety of epigenetic states. a)** Correlation between allele-specific chromatin accessibility (ASCA) and ASE the set of peaks assigned to each chromHMM state. Each peak was assigned to only the state that overlapped the largest proportion of the peak. Significance is indicated by asterisks where *** indicates $p < 0.005$, ** indicates $p < 0.01$, and * indicates $p < 0.05$. **b)** The same as in (a) but with peaks overlapping any promoter-related states (bivalent promoter, flanking promoter, and TSS) removed.

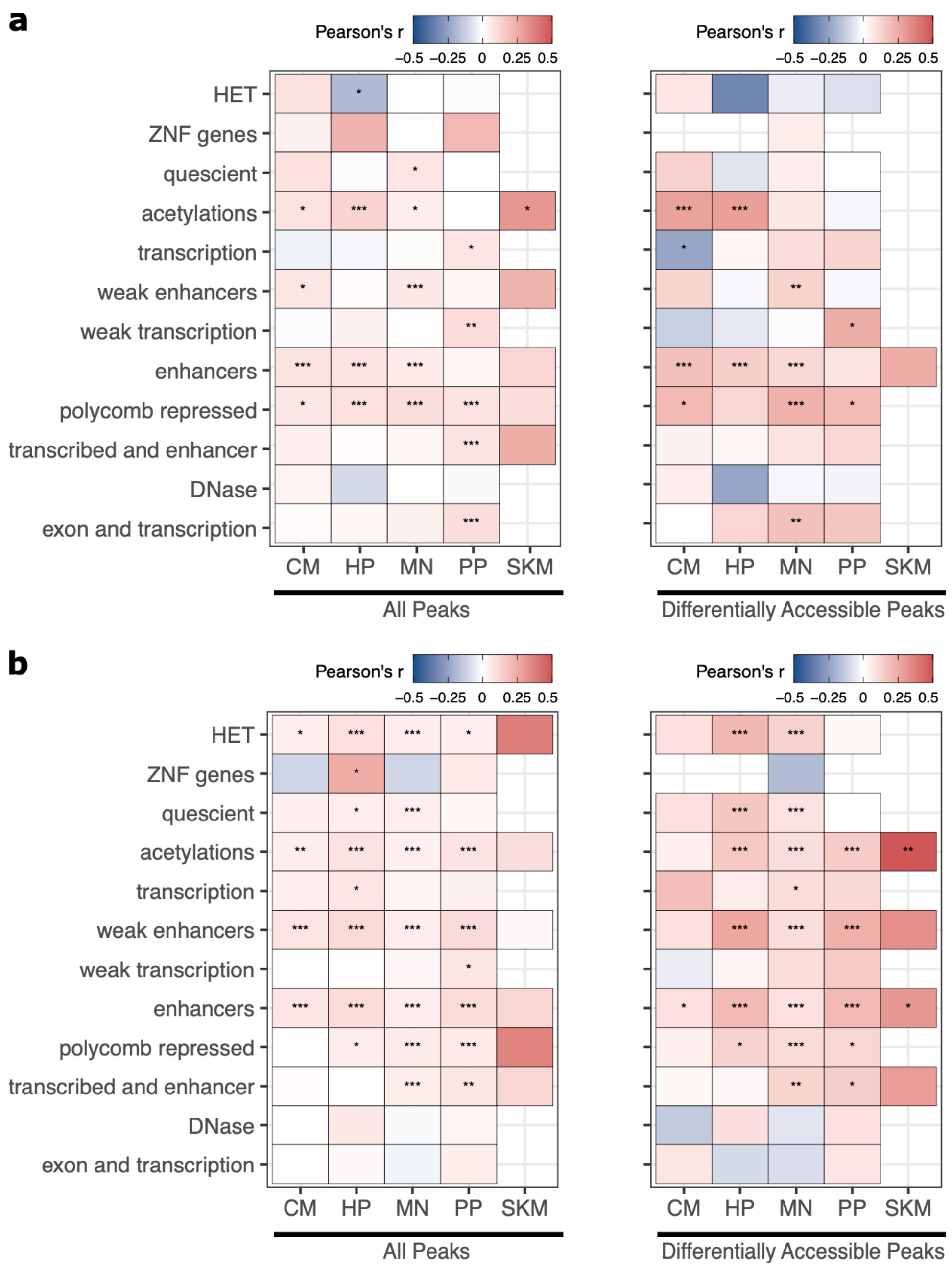


**Supplemental Figure 21**. **ASE and ASCA correlations for TSS-proximal and distal CREs.** Similar to Fig. 3e except peaks were separated into **a)** distance to assigned gene <= 30kb, **b)** distance to assigned gene > 30 kb. Significance is indicated by asterisks where *** indicates $p < 0.005$, ** indicates $p < 0.01$, and * indicates $p < 0.05$.

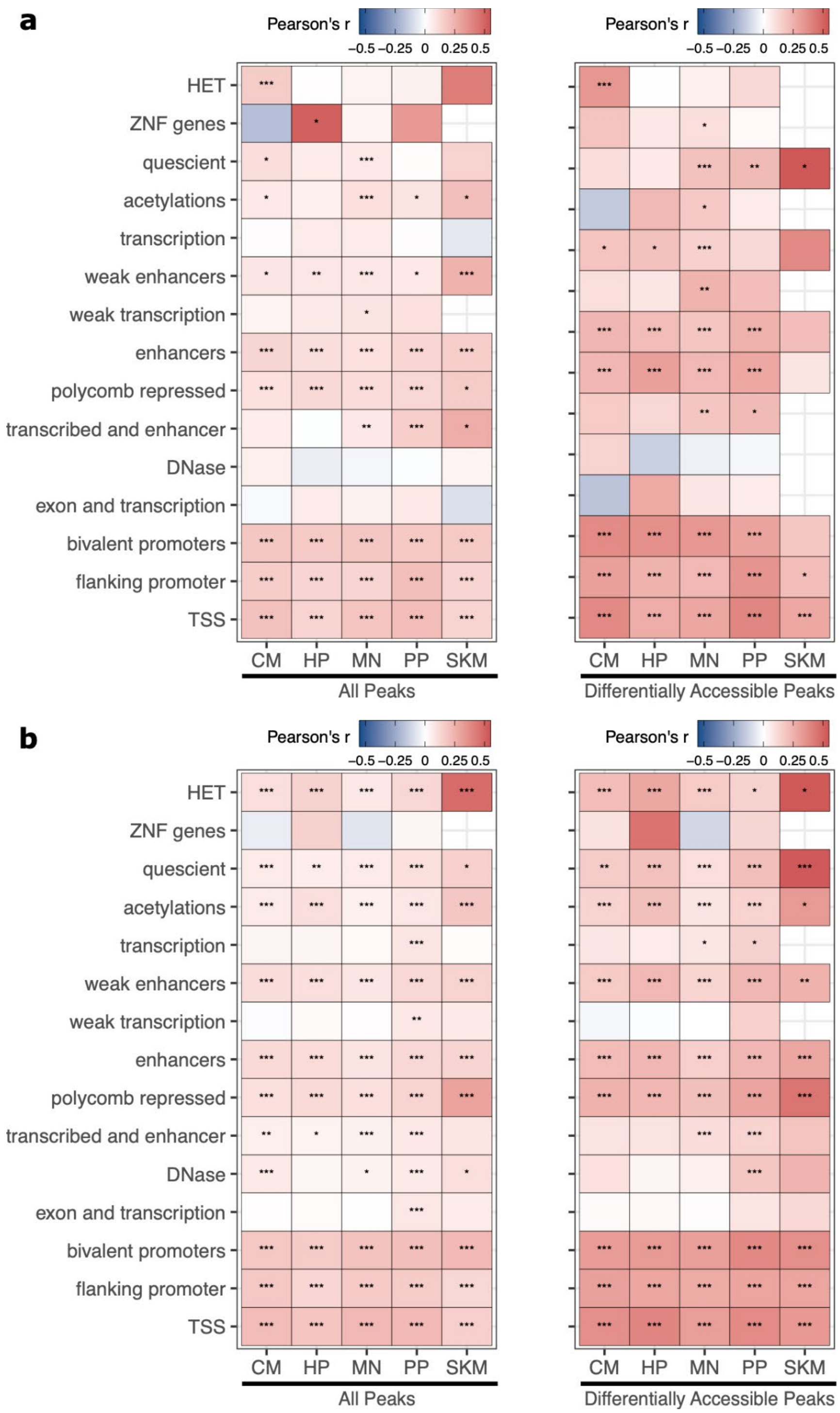


**Supplemental Figure 22**. **ASE and ASCA correlation for cell type-specific and ubiquitously expressed genes.** Similar to Fig. 3e except genes and corresponding peaks were separated into **a)** cell type-specific genes (see Methods) and **b)** ubiquitously expressed genes. Significance is indicated by asterisks where *** indicates $p < 0.005$, ** indicates $p < 0.01$, and * indicates $p < 0.05$.

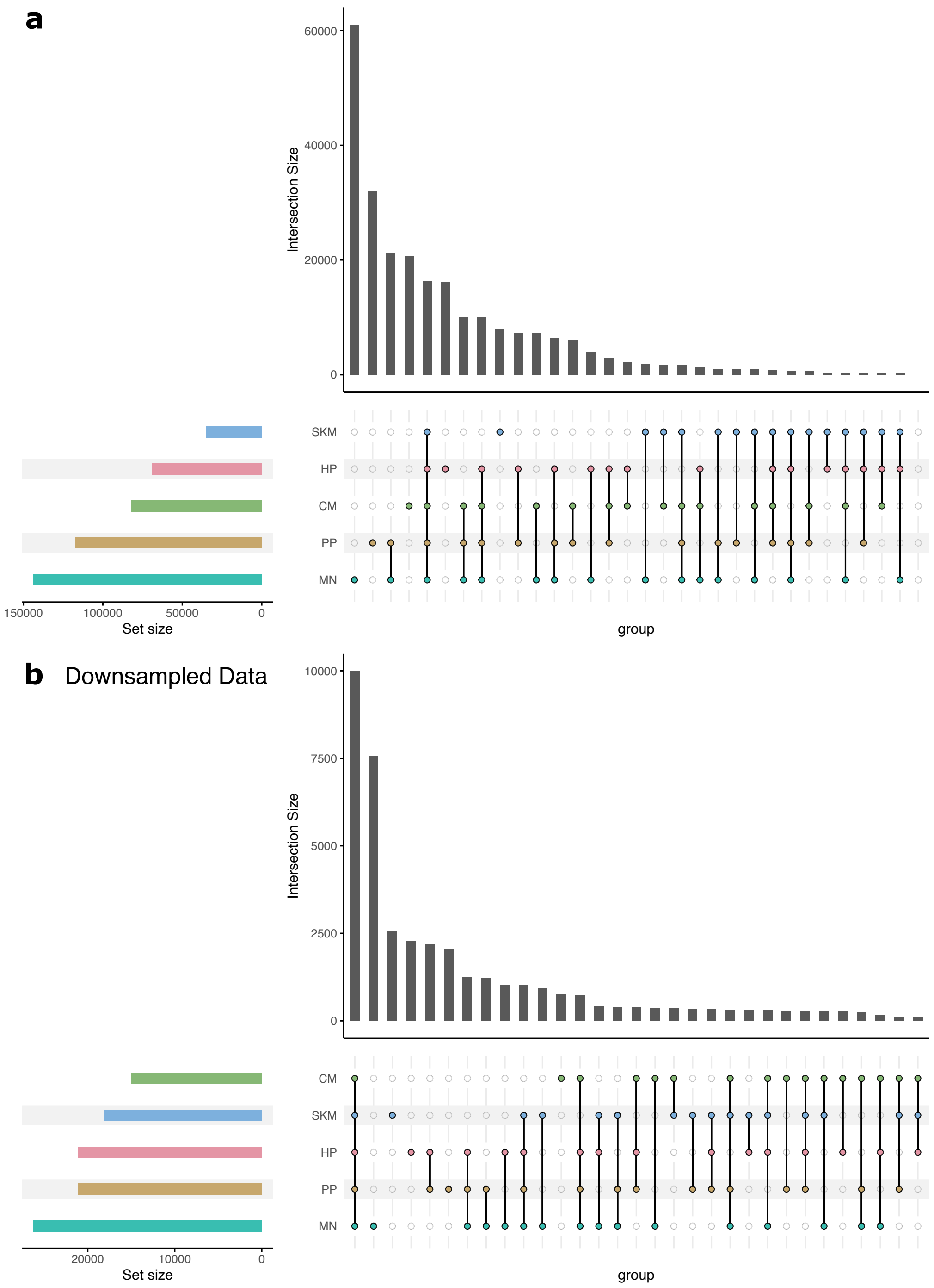


**Supplemental Figure 23. Down-sampling eliminates sequencing depth-induced bias in the number of peaks called in each cell type. a)** Number of peaks called in each cell type and number of common peaks shared among cell types. MN has the largest number of peaks and the largest number of unique peaks primarily because the samples from MN were sequenced the most deeply. **b**) The same as in (a) using a down-sampled dataset so that all cell types have the same read depth. In this case, peaks shared across all cell types make up a much larger proportion of the total number of peaks.

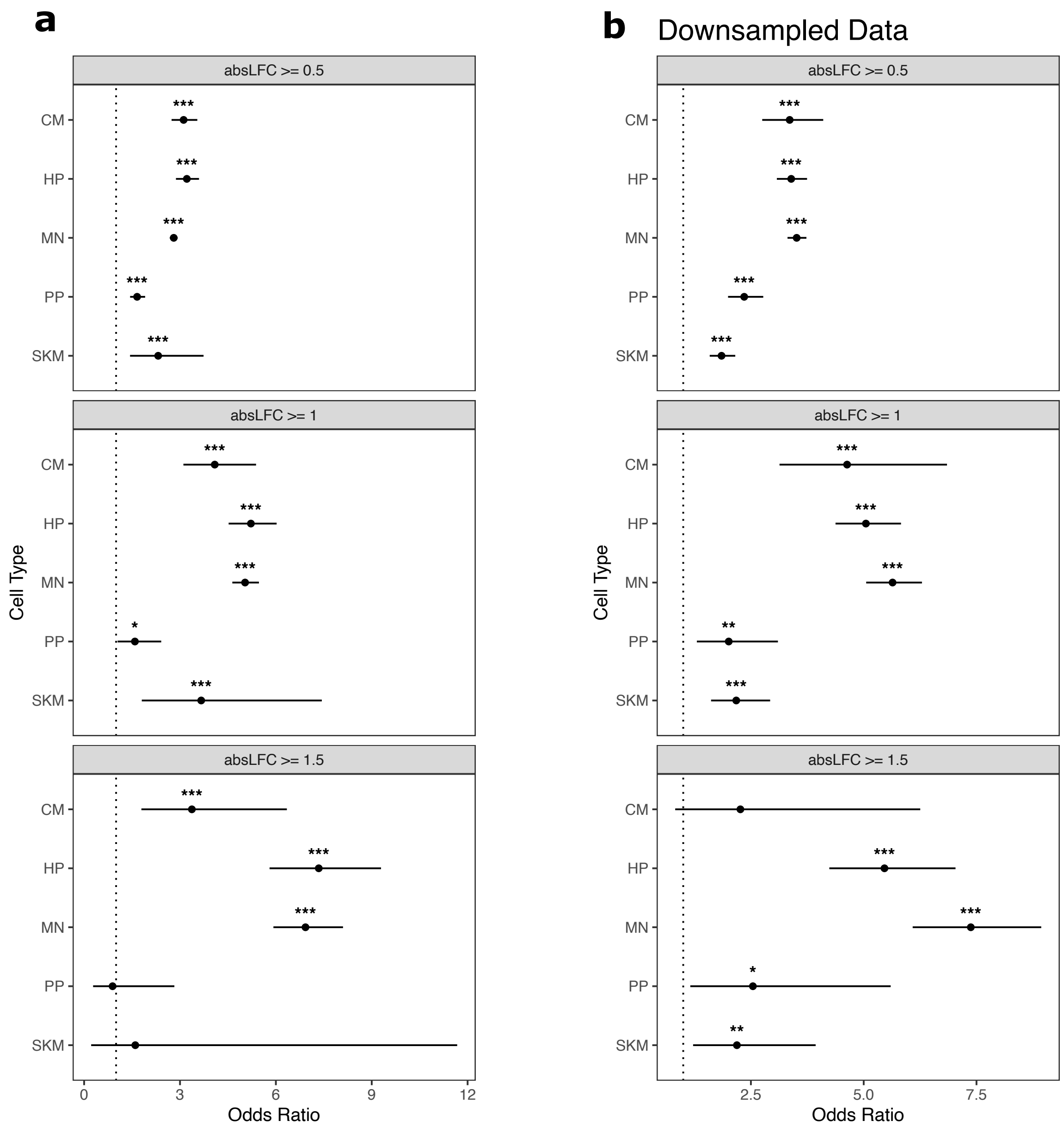


**Supplemental Figure 24. Cell type-specific ATAC-seq peaks are enriched for ASCA across log fold-change cutoffs. A)** Testing if cell type-specific peaks are enriched for peaks showing ASCA. Cell type-specific peaks were defined as peaks only called in that cell type, and peaks showing ASCA are defined using different absolute log fold-change cutoffs. Significance (using the Wald test) is indicated by asterisks where *** indicates $p < 0.005$, ** indicates $p < 0.01$, and * indicates $p < 0.05$. **b)** The same as in (a) but with down-sampled data.

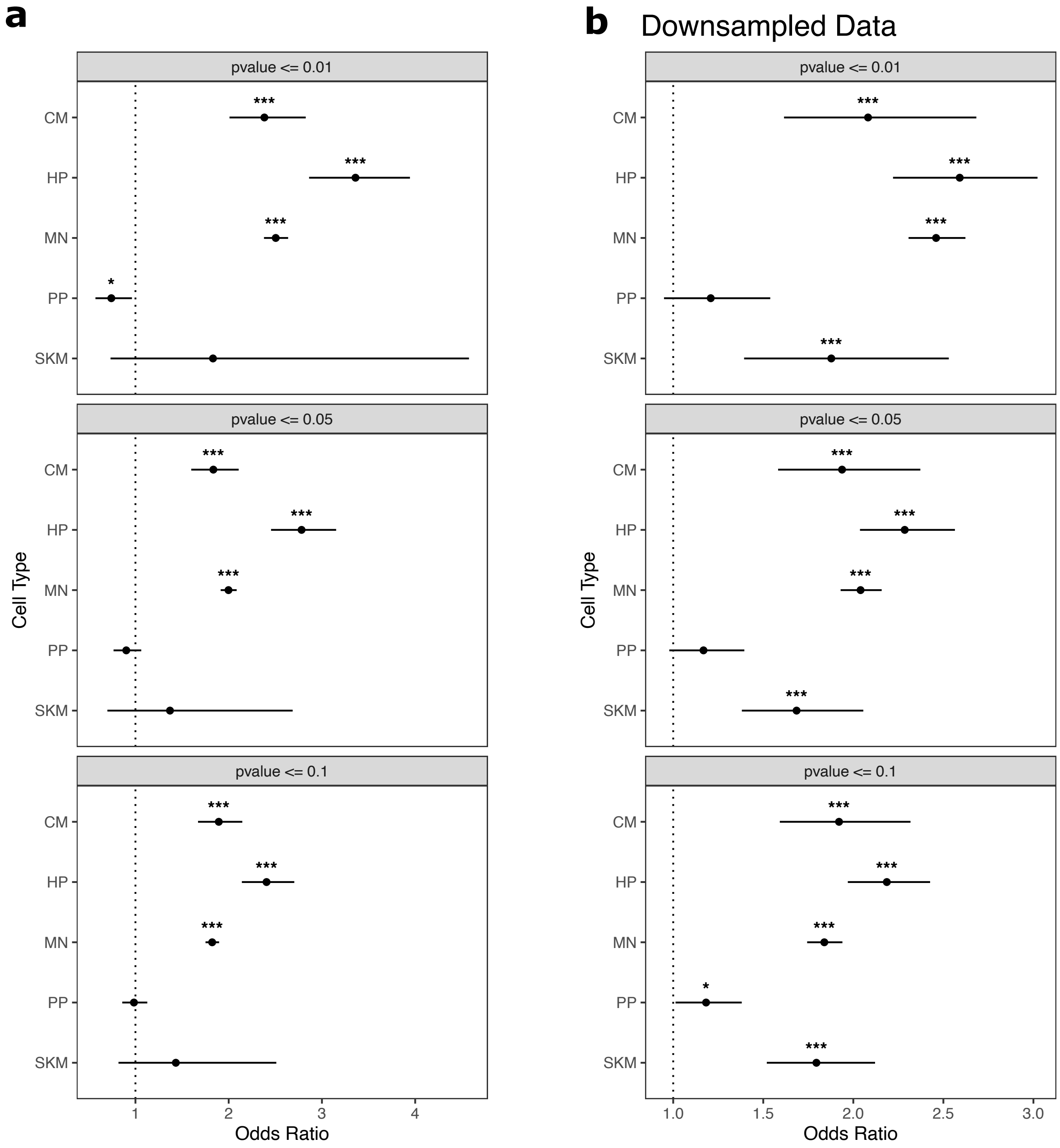


**Supplemental Figure 25. Cell type-specific ATAC-seq peaks are generally enriched for ASCA across p-value cutoffs. a)** Testing if cell type-specific peaks are enriched for peaks showing ASCA using p-value cutoffs. Cell type-specific peaks were defined as peaks only called in that cell type and peaks showing ASCA were defined using different binomial test p-value cutoffs. The read counts from the human and chimpanzee alleles are used as input to the binomial test. Significance (using the Wald test) is indicated by asterisks where *** indicates $p < 0.005$, ** indicates $p < 0.01$, and * indicates $p < 0.05$. **b)** The same is in (a) but with down-sampled data.

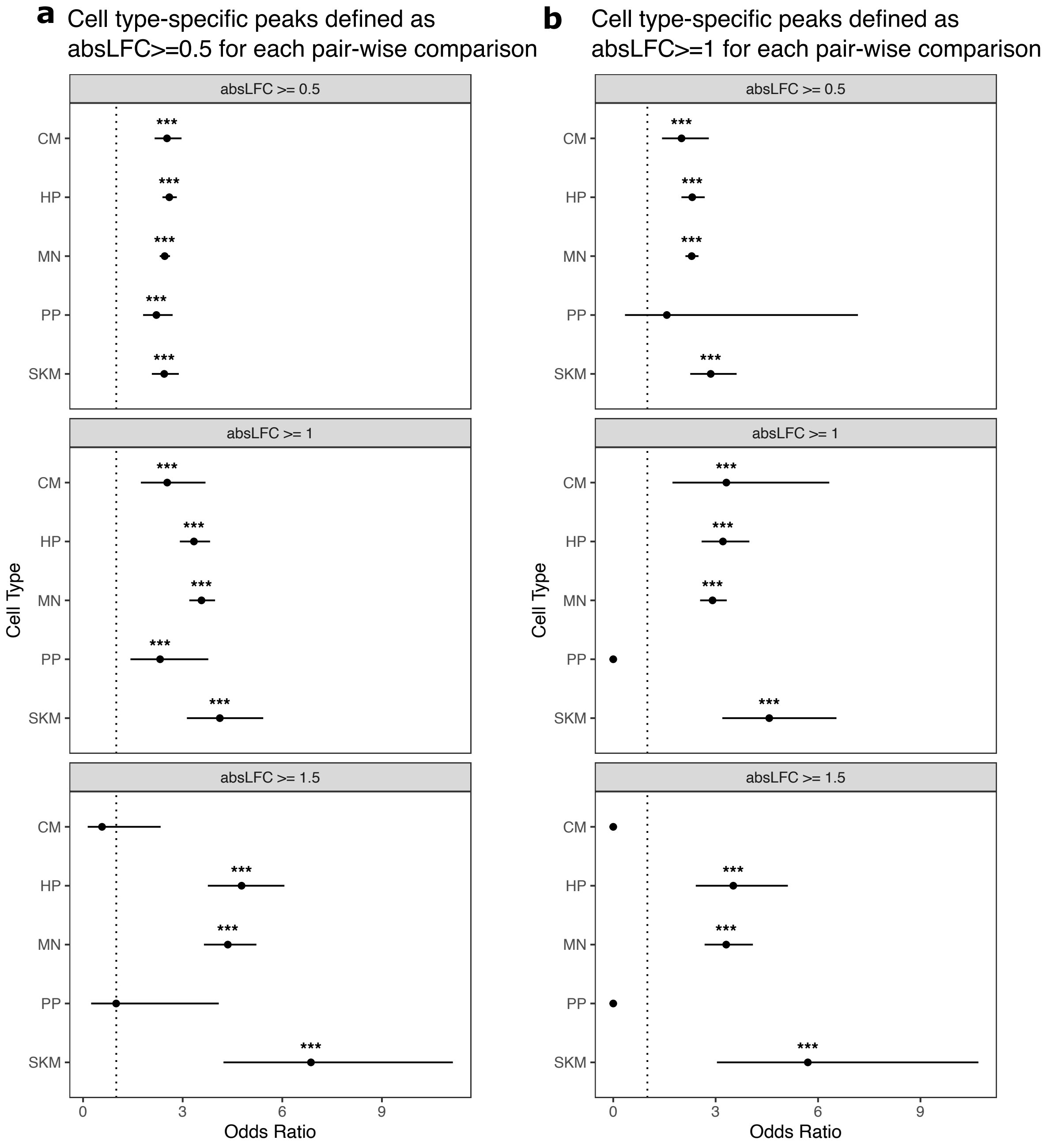


**Supplemental Figure 26. Cell type-specific ATAC peaks are enriched for ASCA using a broader definition of cell type-specific. a)** We defined cell type-specific peaks more broadly as peaks which are differentially accessible between this cell type and all others (i.e., absolute log fold-change greater than 0.5 across all pairwise comparisons between a cell type and all other cell types). Peaks with significant ASCA are defined using different absolute log fold-change cutoffs. All tests were performed on down-sampled data. Significance (using the Wald test) is indicated by asterisks where *** indicates $p < 0.005$, ** indicates $p < 0.01$, and * indicates $p < 0.05$. **b)** The same as in (a) but using an absolute log fold-change cutoff of 1 to define cell type-specific ATAC peaks.

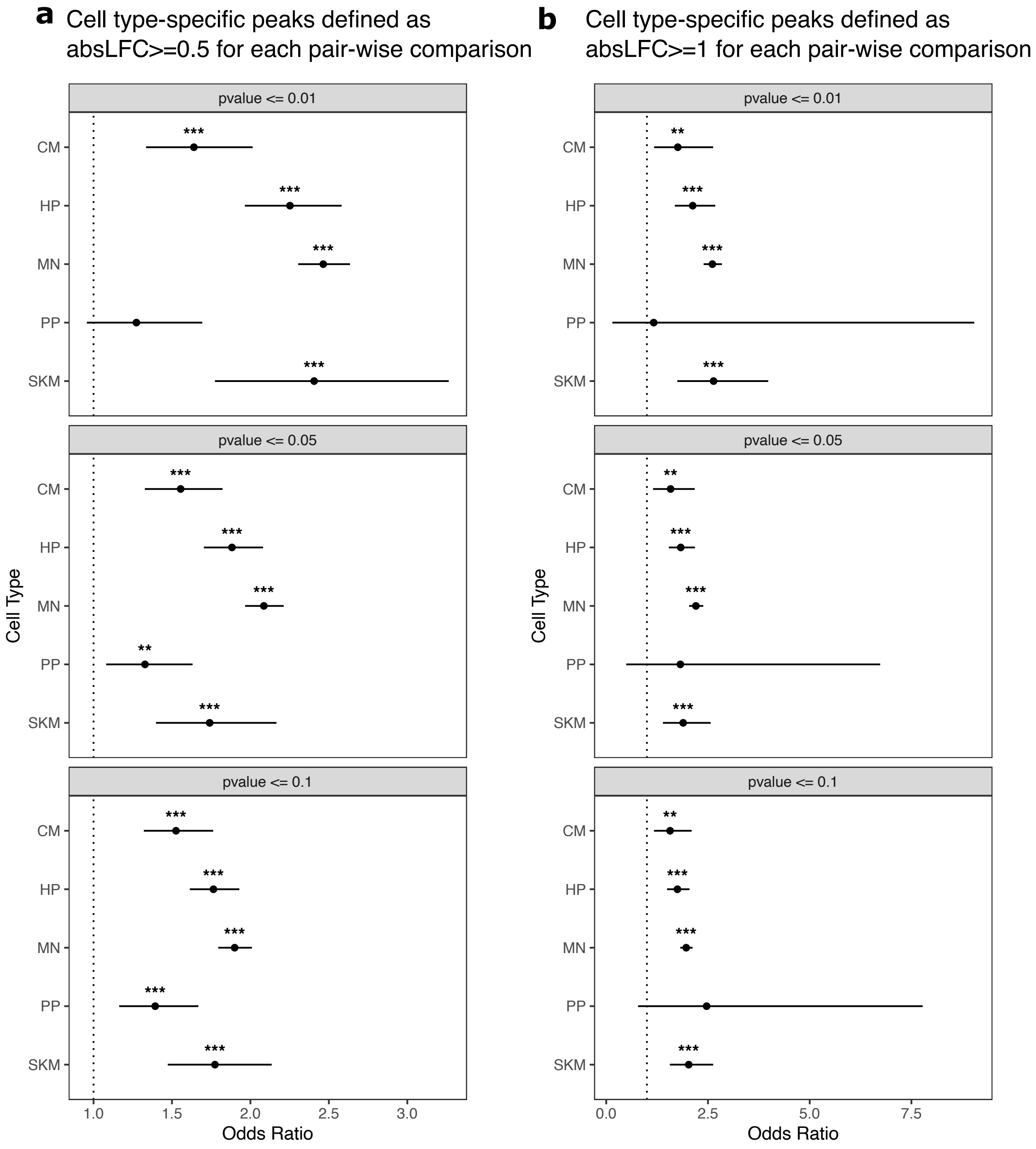


**Supplemental Figure 27. Cell type-specific ATAC peaks are enriched for ASCA using a broader definition of cell type-specific. a)** We defined cell type-specific peaks more broadly as peaks which are differentially accessible between this cell type and all others (i.e., absolute log fold-change greater than 0.5 across all pairwise comparisons between a cell type and all other cell types). Peaks with significant ASCA are defined using different p-value cutoffs from the binomial test applied to human and chimpanzee allelic counts. All tests were performed on down-sampled data. Significance (using the Wald test) is indicated by asterisks where *** indicates $p < 0.005$, ** indicates $p < 0.01$, and * indicates $p < 0.05$. **b)** The same as in (a) but using an absolute log fold-change cutoff of 1 to define cell type-specific ATAC peaks.

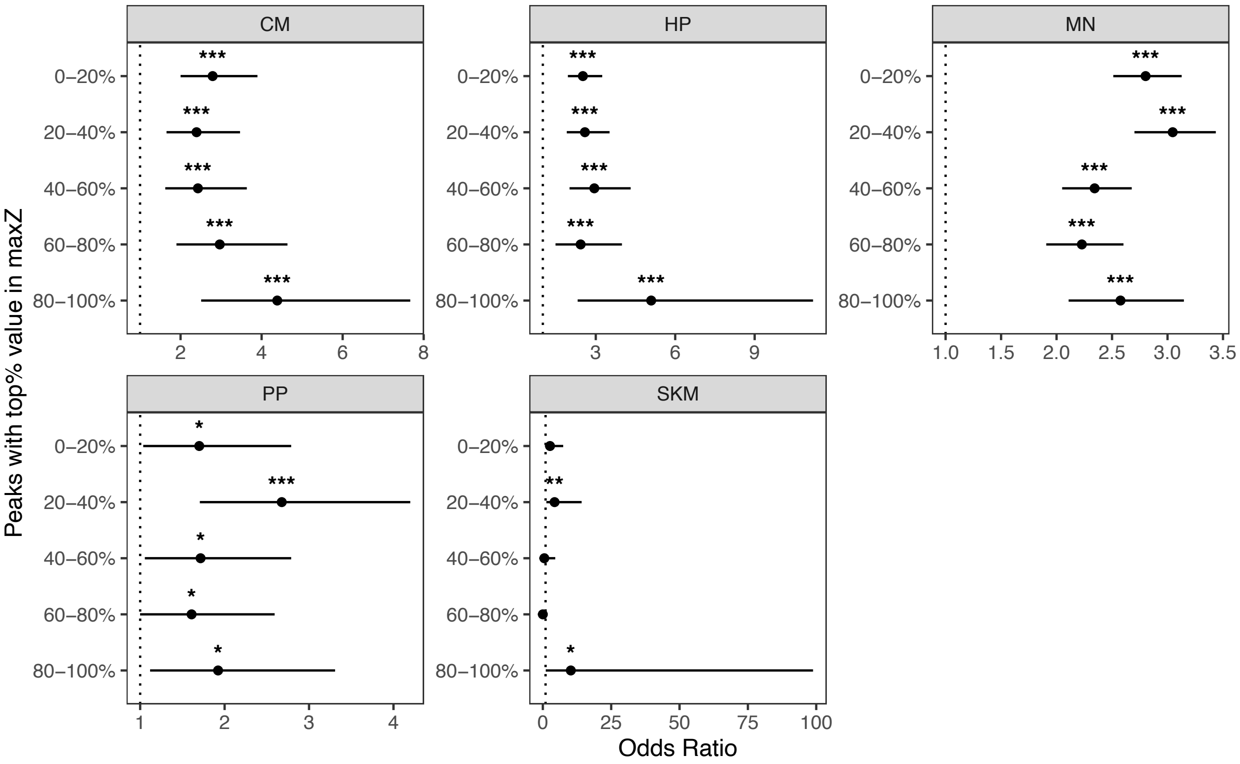


**Supplemental Figure 28. Cell type-specific ATAC peaks are generally enriched for ASCA regardless of sequence constraint.** Testing if cell type-specific peaks are enriched for peaks showing ASCA when stratifying by conservation of genomic regions. Genes were split into equal size bins based on the maximum z-score value (maxZ), a metric for constraint based on comparing the observed number of mutations in a region to the expected number of mutations in a large set of whole genome sequencing samples from humans. Peaks in the 0-20% bin are the least constrained whereas peaks in the 80-100% bin are the most constrained. Cell type-specific peaks were defined as peaks only called in that cell type, and differentially accessible peaks are defined as those with an absolute log fold-change greater than or equal to 0.5. Significance (using the Wald test) is indicated by asterisks where *** indicates $p < 0.005$, ** indicates $p < 0.01$, and * indicates $p < 0.05$.

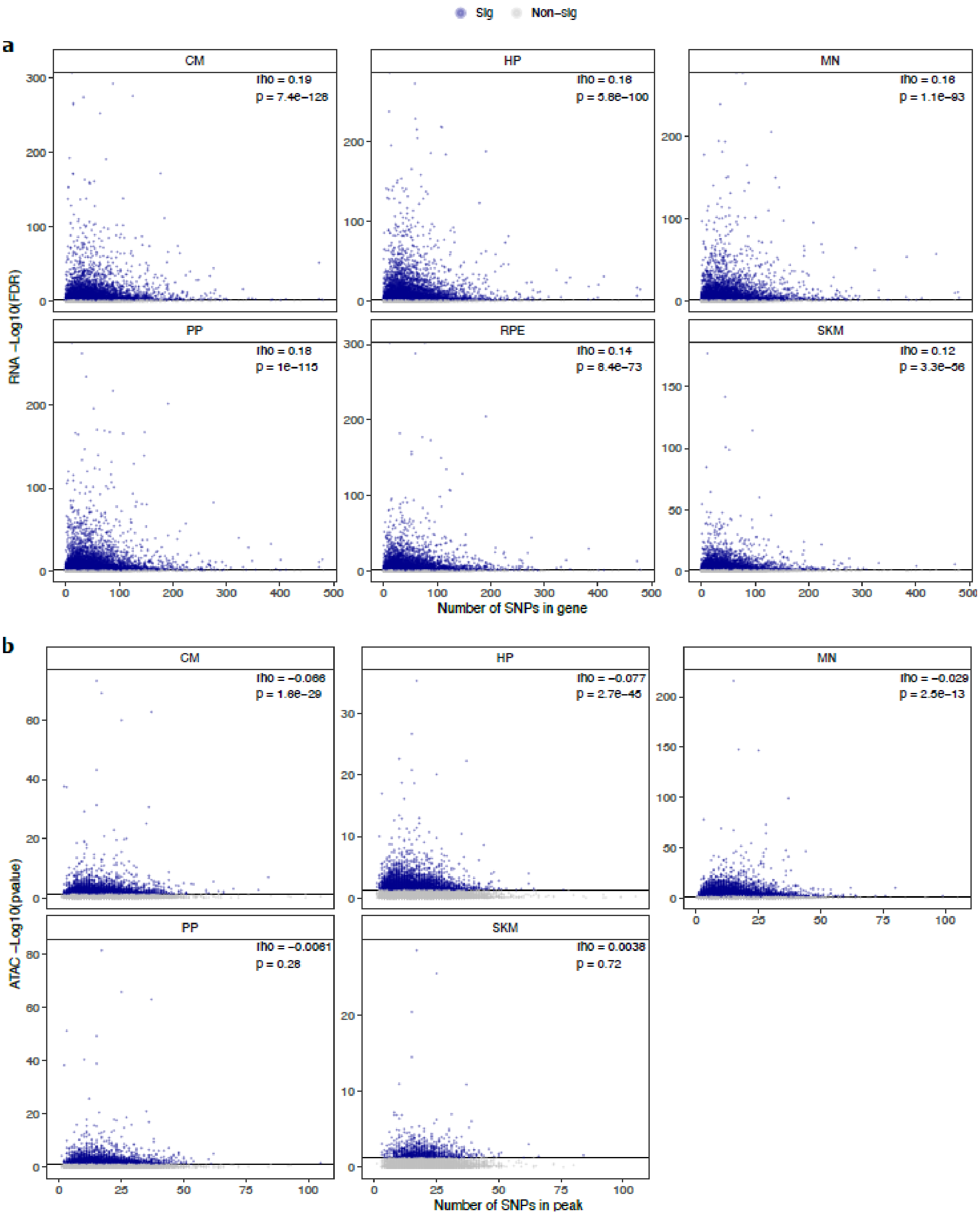


**Supplemental Figure 29. Relationship between number of SNPs and** -$\log_{10}$(FDR) in a) ASE and -$\log_{10}$(p-value) b) ASCA. These scatter plots show the relationship between the number of SNPs in a gene or peak and the -$\log_{10}$(FDR) for ASE or ASCA. Genes with significant ASE (FDR < 0.05) and peaks with significant ASCA (binomial p-value < 0.05) were annotated as blue dots, and all other genes and peaks were annotated as gray dots. All genes in each cell type in RNA-seq are shown. For clarity, the few outlier peaks with more than 200 SNPs are excluded from these plots.

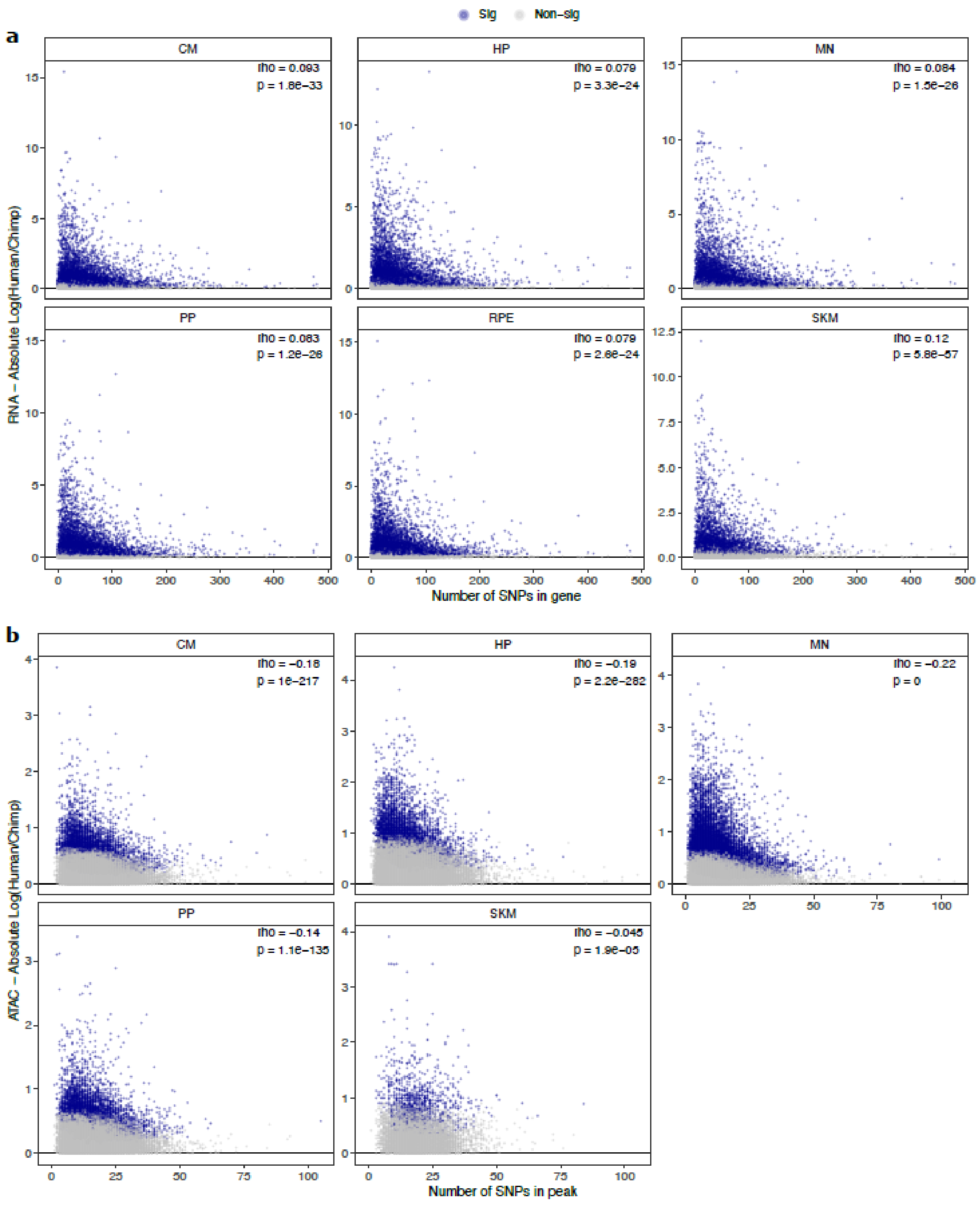


**Supplementary Figure 30. Relationship between number of SNPs and absolute $\log_2$ fold-change** in a) ASE and b) ASCA. These scatter plots show the relationship between the number of SNPs in a gene or peak and the estimated absolute $\log_2$ fold-change for ASE or ASCA. Genes with significant ASE (FDR < 0.05) and peaks with significant ASCA (binomial p-value < 0.05) were annotated as blue dots, and all other genes and peaks were annotated as gray dots. All genes in each cell type in RNA-seq are shown. For clarity, the few outlier peaks with more than 200 SNPs are excluded from these plots.

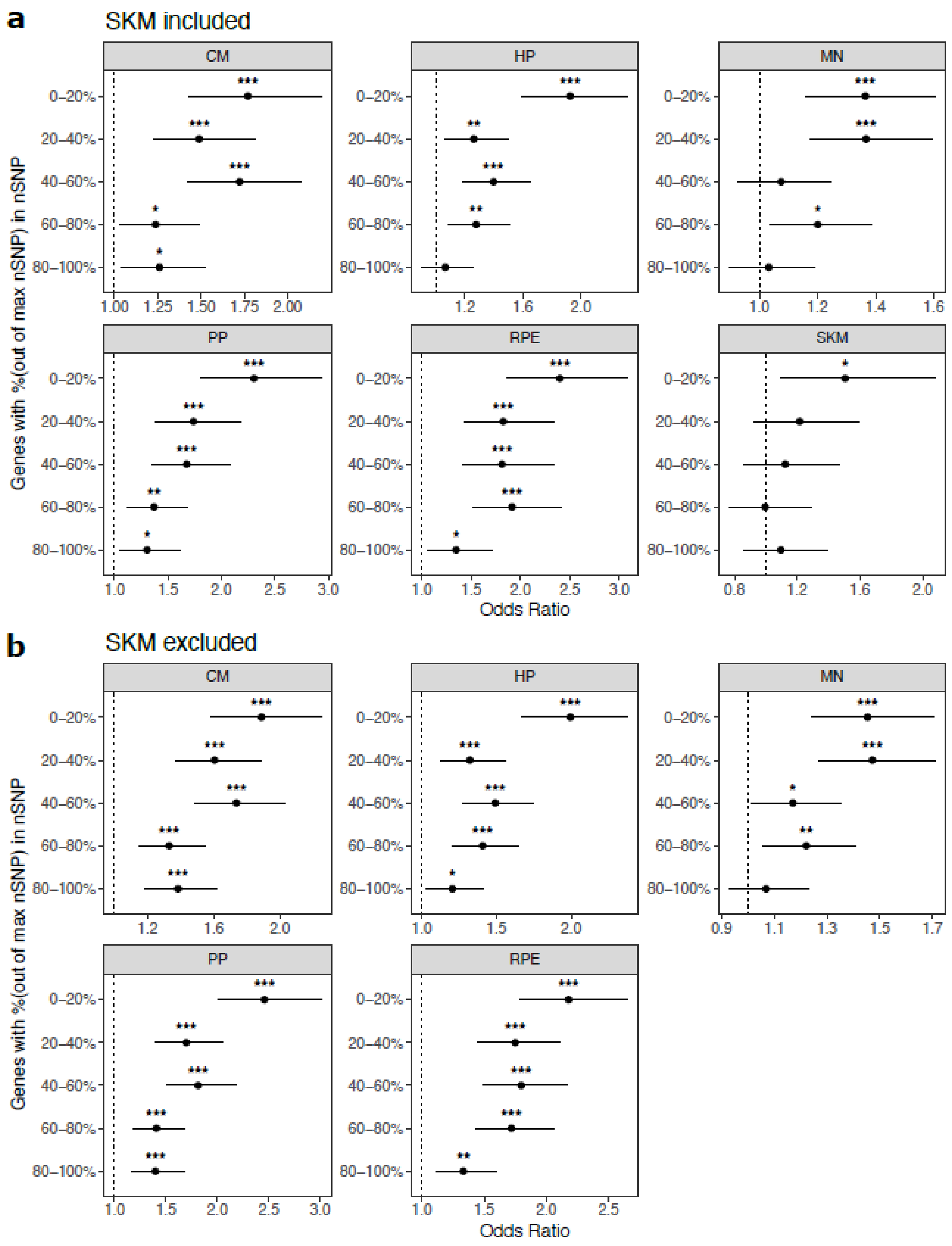


**Supplemental Figure 31. Cell type-specifically expressed genes are enriched for genes with ASE when stratifying by the number of SNPs per gene. a)** Results when SKM is included. Genes were put into five bins with an equal number of genes in each bin. Genes with the fewest SNPs are in the 0-20% bin and genes with the most SNPs are in the 80-100% bin. Significance (using the Wald test) is indicated by asterisks where *** indicates $p < 0.005$, ** indicates $p < 0.01$, and * indicates $p < 0.05$. **b)** The same as in (a) but excluding SKM.

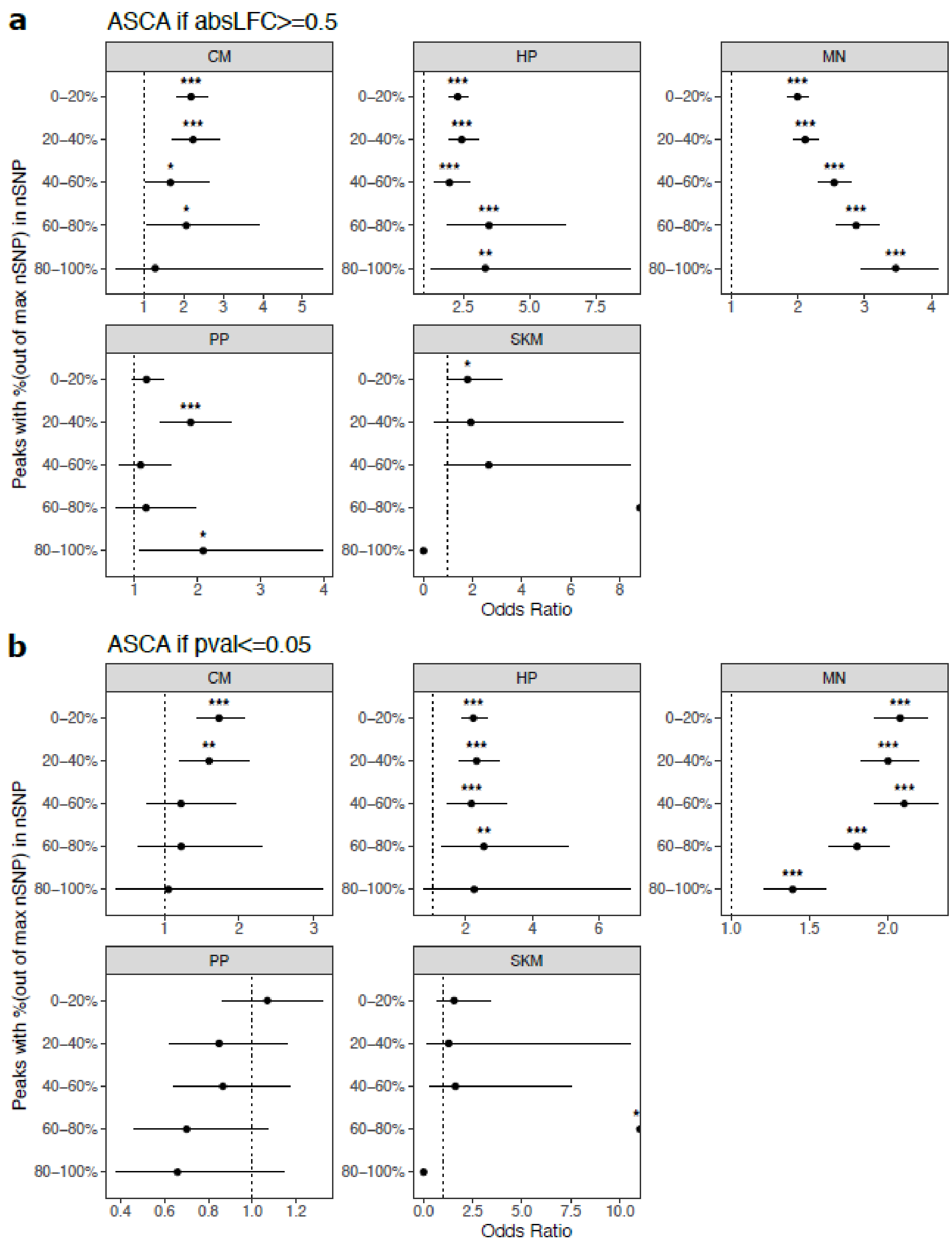


**Supplemental Figure 32. Cell type-specific peaks are enriched for ASCA when stratifying by the number of SNPs per peak. a)** Peaks with an absolute $\log_2$ fold-change greater than or equal to 0.5 were called as having ASCA. Peaks were put into five bins with an equal number of peaks in each bin. Peaks with the fewest SNPs are in the 0-20% bin and genes with the most SNPs are in the 80-100% bin. Significance (using the Wald test) is indicated by asterisks where *** indicates $p < 0.005$, ** indicates $p < 0.01$, and * indicates $p < 0.05$. **b)** The same as in (a) but peaks with a binomial p-value less than or equal to 0.05 were called as having ASCA.

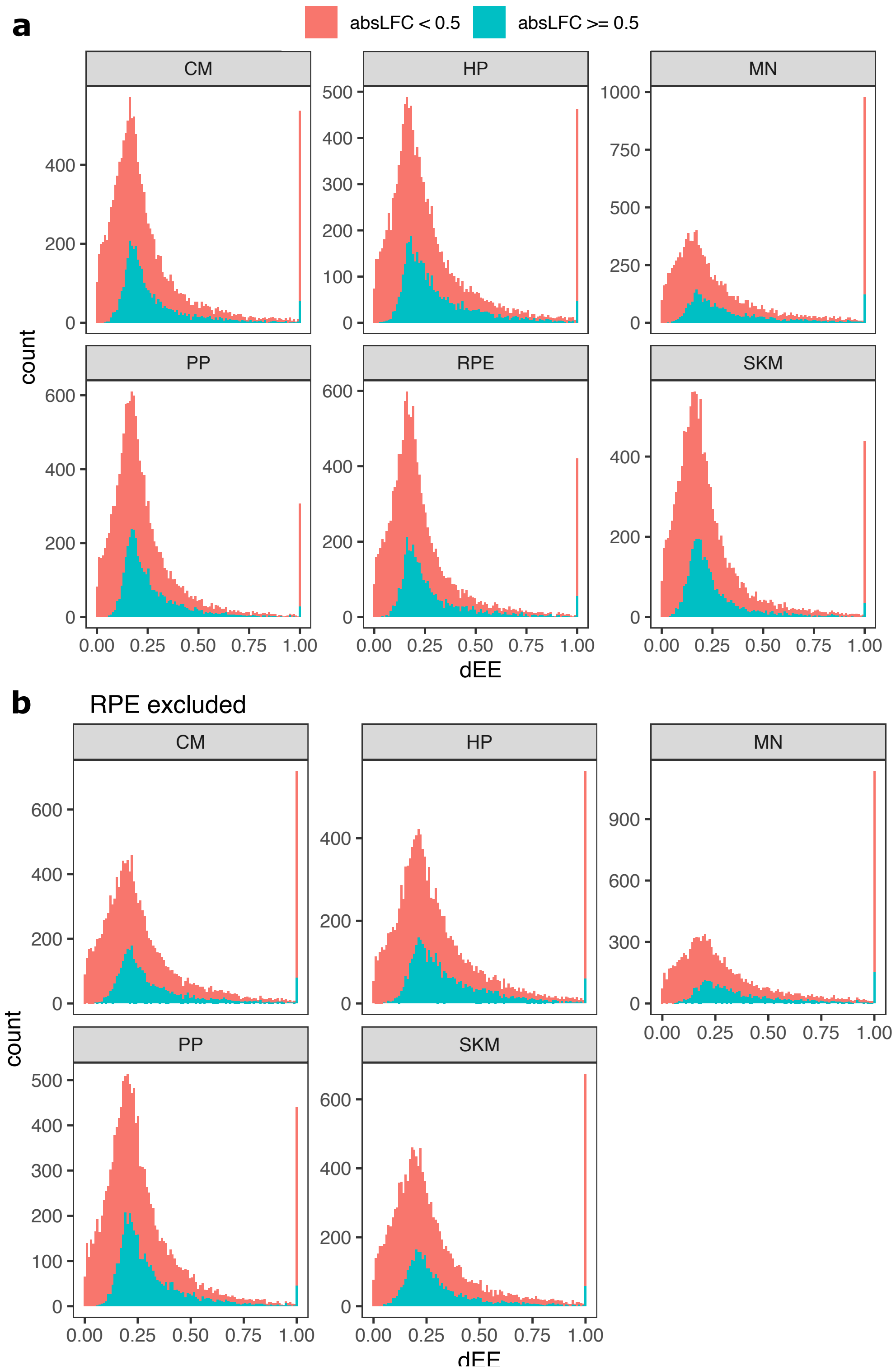


**Supplemental Figure 33. Distribution of dEE across cell types. a)** Histogram of differential expression enrichment (dEE) computed on ASE ratios for each cell type. The red color indicates inclusion of all genes whereas the blue color indicates inclusion of only genes with an absolute log fold-change greater than 0.5. **b)** The same as in (a) but excluding RPE to more closely match the ATAC-seq dataset.

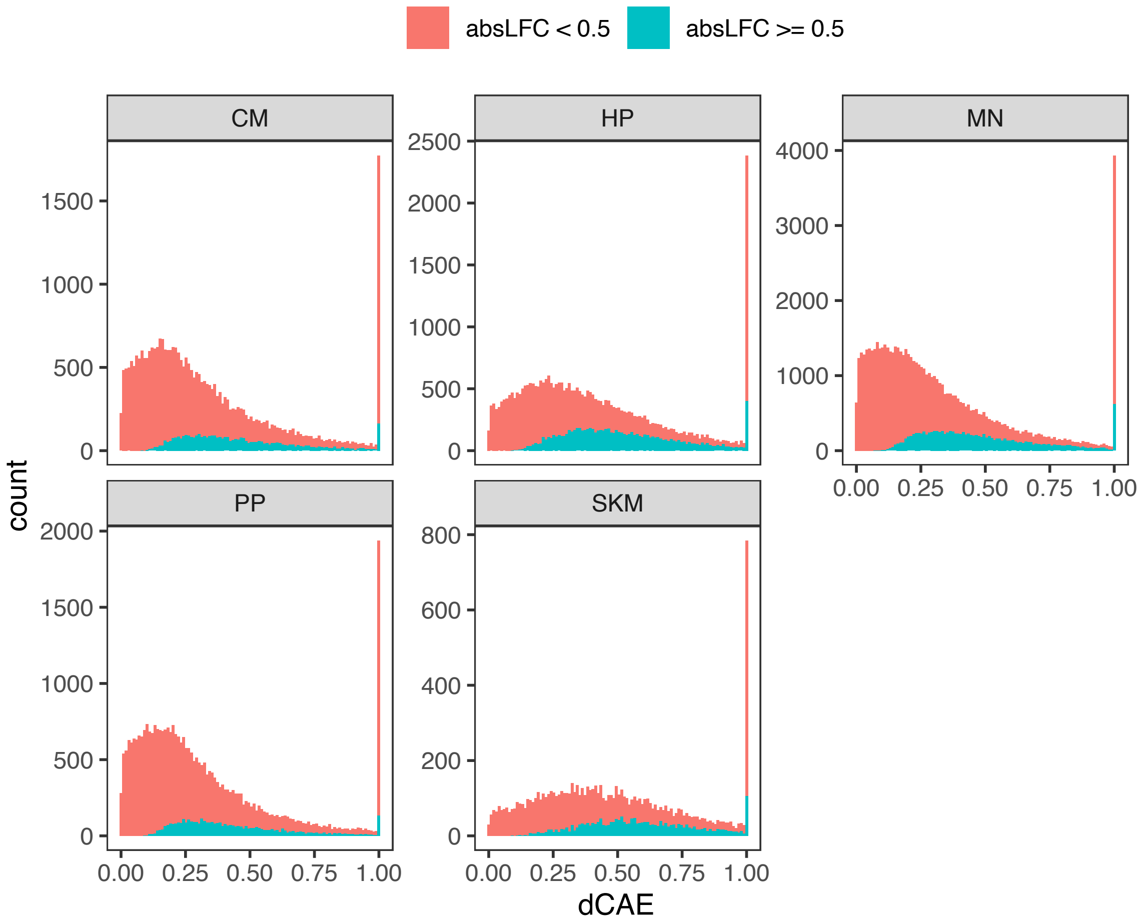


**Supplemental Figure 34. Distribution of dCAE across cell types. a)** Histogram of differential chromatin accessibility enrichment (dCAE) computed on ASCA ratios for each cell type. The red color indicates inclusion of all genes whereas the blue color indicates inclusion of only genes with an absolute log fold-change greater than 0.5.

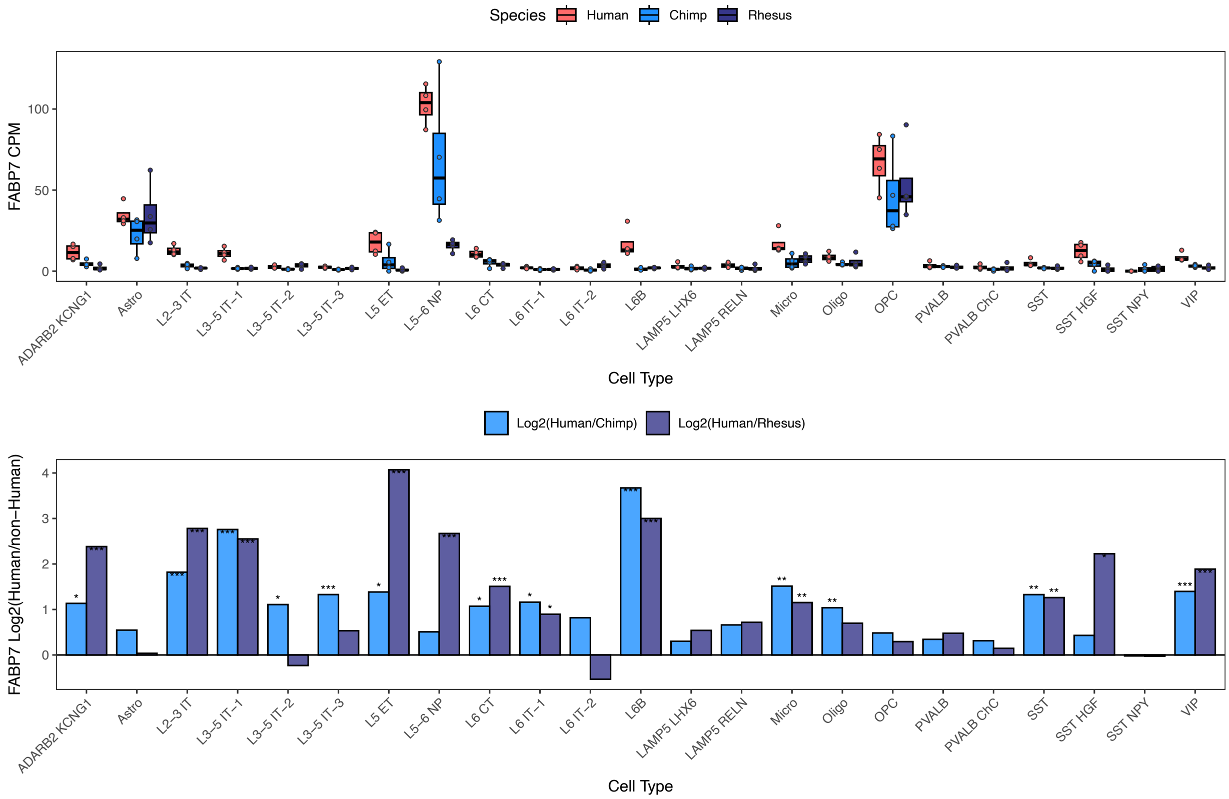


**Supplemental Figure 35. Differential *FABP7* expression across cortical cell types.** *FABP7* expression in the human, chimpanzee and rhesus snRNA-seq dataset from the dorsolateral prefrontal cortex collected by Ma et al. **a)** CPM and **b)** $\log_2$ fold-change between human and non-human (chimpanzee or rhesus) are shown. Significance (using the DESeq2 likelihood ratio test) is indicated by asterisks where *** indicates FDR < 0.005, ** indicates FDR < 0.01, and * indicates FDR < 0.05. Astro: astrocytes, Micro: microglia, Oligo: oligodendrocytes, OPC: oligodendrocyte progenitor cells. ADARB2 KCNG1, LAMP5 LHX6, LAMP5 RELN, PVALB, PVALB ChC, SST, SST HGF, SST NPY, and VIP are inhibitory interneuron cell types. IT (intratelencephalic), ET (extratelencephalic), NP (near-projecting), and CT (corticothalamic) neuronal types are excitatory projection neurons. As an example, L2-3 stands for cortical layers 2-3 and L5-6 stands for cortical layers 5-6.

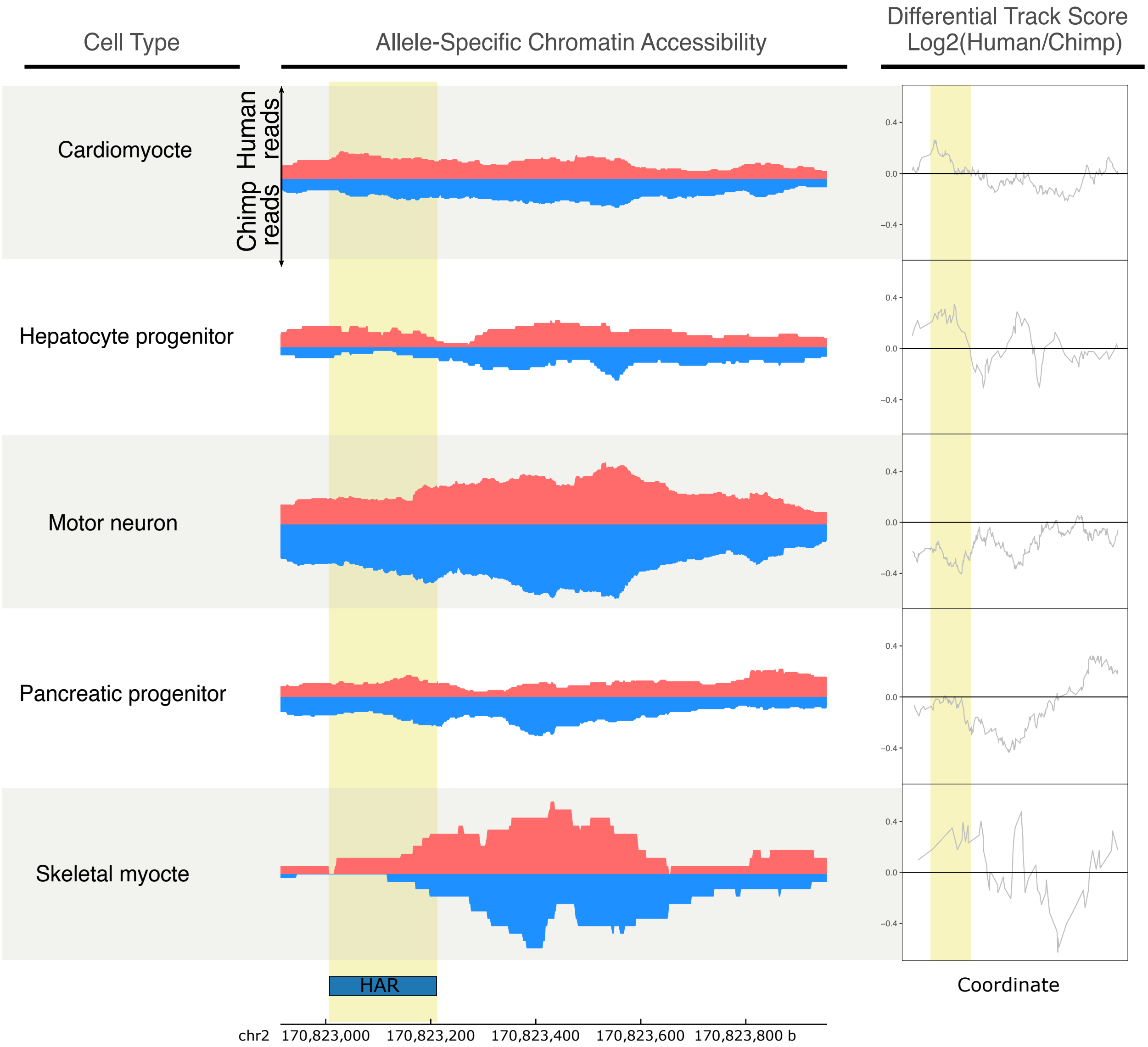


**Supplemental Figure 36: A HAR near the TSS of *GAD1* is chimpanzee-biased only in motor neurons.** Allelic ATAC-seq tracks for a peak near the *GAD1* TSS that contains a HAR (Fig. 5a). The human signal is shown in red, and the chimpanzee signal is shown in blue. Differential track scores ($\log_2$(human/chimp)) were computed by position along the genomic coordinates and are shown on the right. The HAR region is highlighted in yellow. Motor neuron-specific chimpanzee biased accessibility was observed in the HAR region.

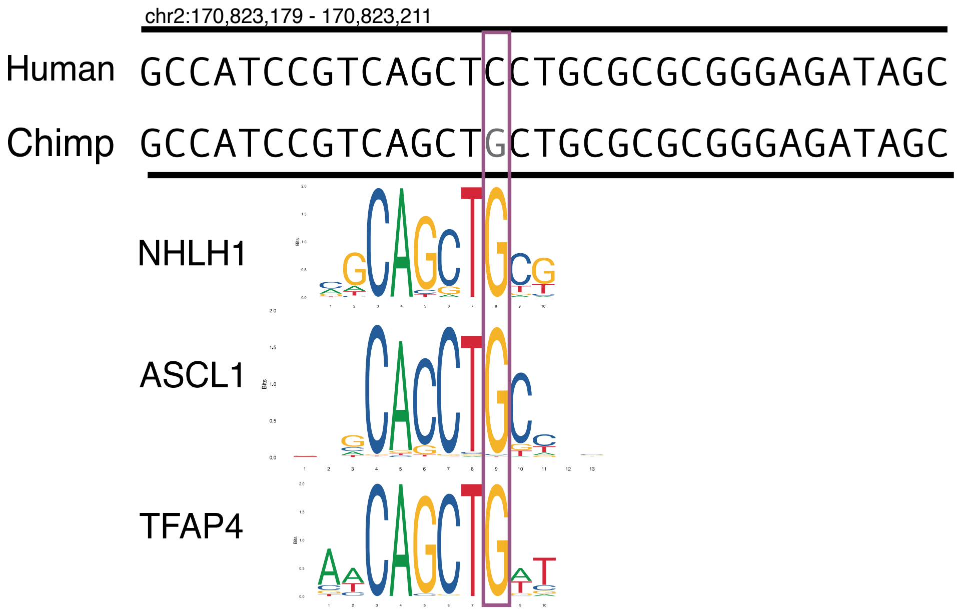


**Supplemental Figure 37. A human-specific SNP disrupts a potential Ascl1 binding site.** The potential effects of the human-chimpanzee difference at hg38 position chr2:170,823,193 (C in humans and G in chimpanzees) on transcription factor binding sites are shown.

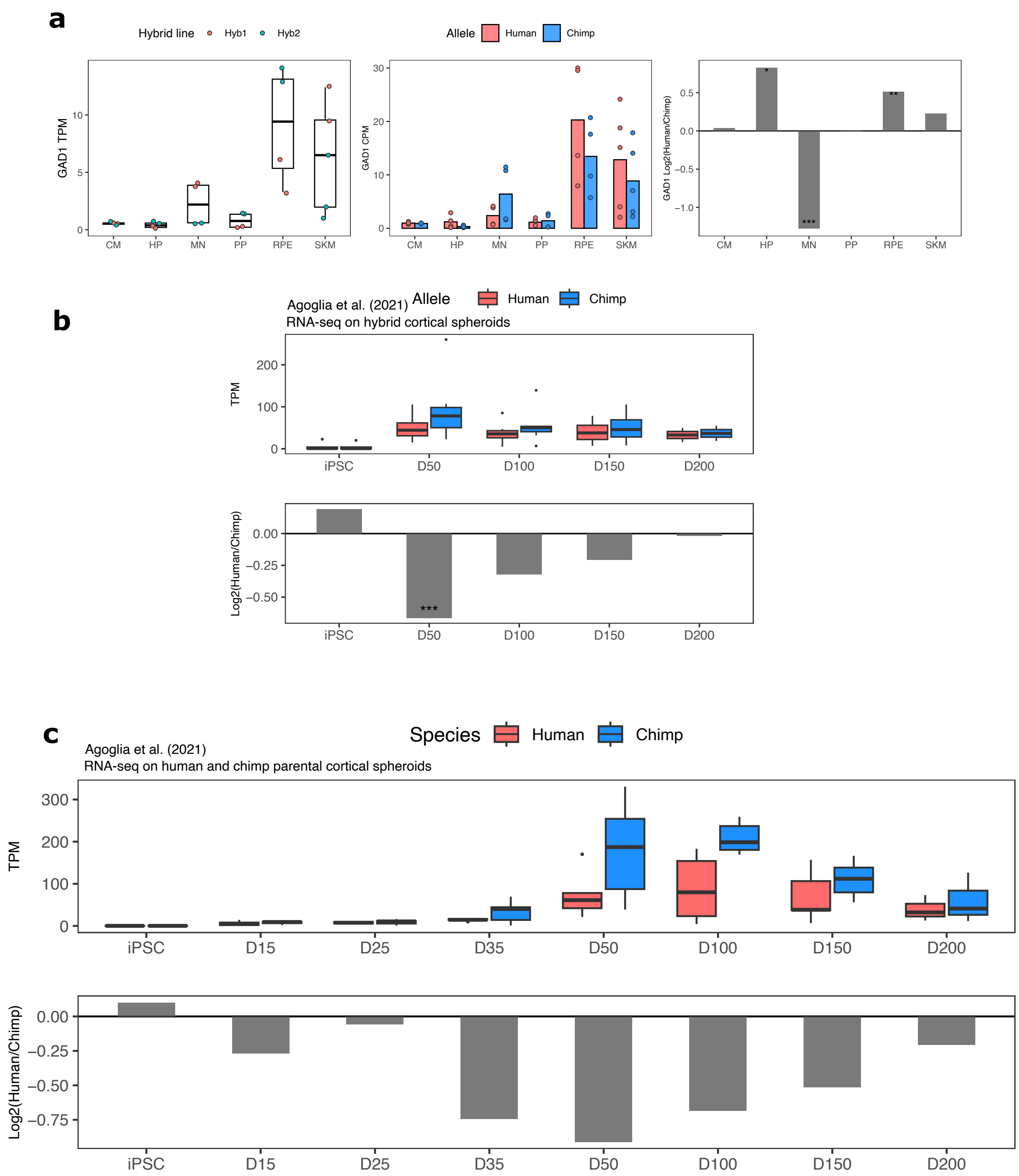


**Supplemental Figure 38.** ***GAD1*** **ASE and differential expression across cell types and in cortical organoids. a)** *GAD1* TPM, ASE and DESeq2 log fold-change in all cell types in our dataset. Significance (using the DESeq2 likelihood ratio test) is indicated by asterisks where *** indicates FDR < 0.005, ** indicates FDR < 0.01, and * indicates FDR < 0.05. **b)** *GAD1* ASE in hybrid cortical organoids at different time points. Allelic TPM is shown in the top plot and DESeq2 log fold-change is shown in the bottom plot. **c)** *GAD1* gene expression in human and chimpanzee parental cortical organoids. Allelic TPM is shown in the top plot and DESeq2 log fold-change is shown in the bottom plot.

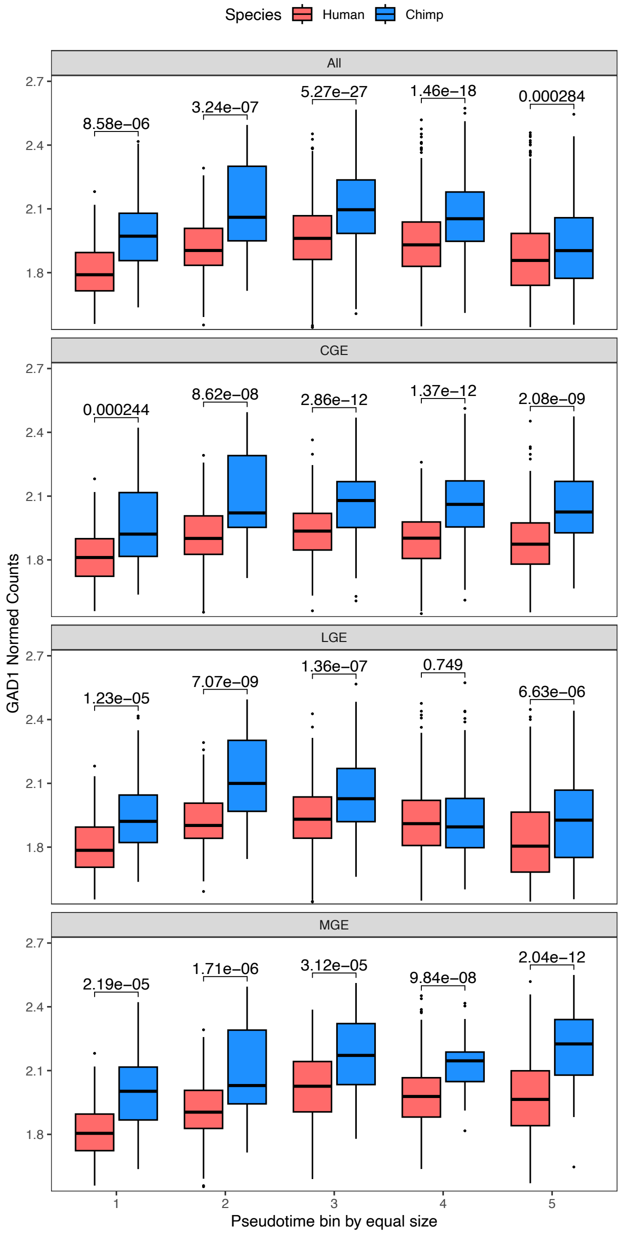


**Supplemental Figure 39. *GAD1* expression across GABAergic neuron differentiation in human and chimpanzee cortical organoids (Kanton et al. 2019).** The top panel shows the results when including all GABAergic neurons regardless of lineage. This closely matches the results when restricting to cells likely to represent the CGE (caudal ganglionic eminence), LGE (lateral ganglionic eminence), or MGE (medial ganglionic eminence) indicating that this difference is likely to affect all forebrain GABAergic lineages. Pseudotime was computed and cells were split into five equally sized bins with bin 1 having the lowest pseudotime value and bin 5 the highest. As no subclusters corresponding to the CGE, MGE, or LGE could be discerned in the progenitor cluster, all progenitors were included in each lineage and fall only into the first two pseudotime bins in each plot. Wilcoxon tests were performed on the non-zero counts and p-values are reported in each pseudotime bin on the figure.

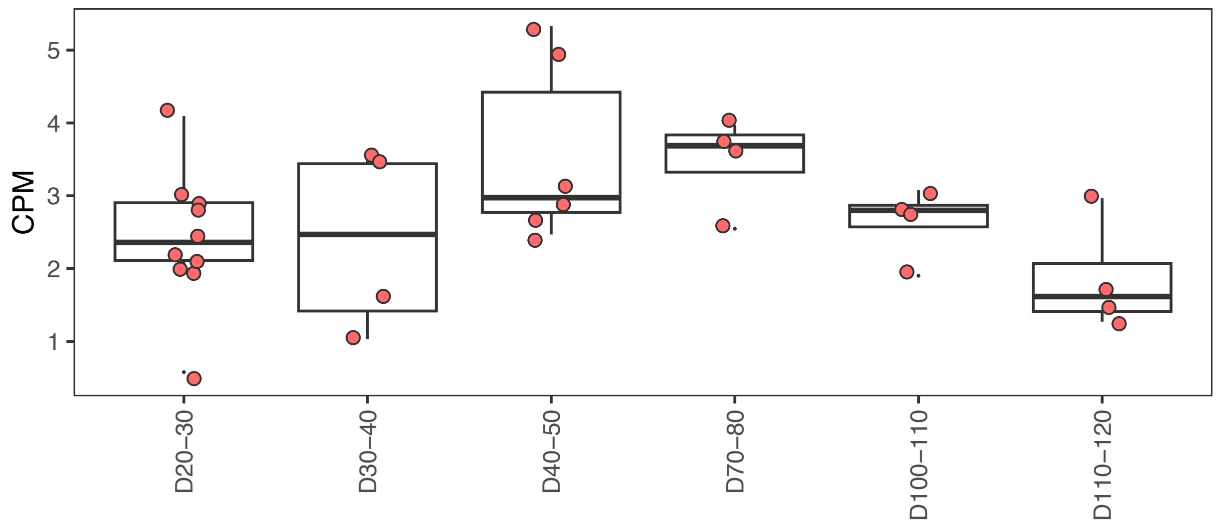


**Supplemental Figure 40. Chromatin accessibility of the HAR near the *GAD1* TSS across human striatal organoid development.** The peak containing the HAR near *GAD1* shows a peak in accessibility during development in human from ATAC-seq on human striatal organoids from Trevino et al.
