## Additional file 1 for "Cell type-specific *cis*-regulatory divergence in gene expression and chromatin accessibility revealed by human-chimpanzee hybrid cells"

Before continuing with the analysis, we tested whether the differentiations were successful and contained primarily our target cell types. The very low expression of *NANOG*, a marker for pluripotency, across all differentiations indicates that the samples contain very few iPSCs (Agoglia et al., 2021). For cardiomyocytes (CM), *NKX2-5*, *MYBPC3*, and *TNNT2* definitively distinguish CM from other heart cell types and their high expression indicates successful differentiations (Burridge et al., 2014). For motor neurons, the high expression of *ELAVL2*, a pan-neuronal marker, indicates a high abundance of neurons in the sample (Mickelsen et al., 2019). The expression of *ISL1* and *OLIG2* further demonstrates that these are motor neurons and not other types of neurons (Maury et al., 2015). For retinal pigment epithelium (RPE), the combined expression of *MITF*, *PAX6*, and *TYRP1* provides strong evidence that the differentiations were successful in producing RPE cells (Sharma et al., 2019). For skeletal muscle, the very high expression of *MYL1*, *MYLPF*, and *MYOG* indicates that these samples contain a high proportion of skeletal muscle cells (Chal et al., 2016). In general, all these populations of cells contain some proportion of progenitors as there is detectable expression of *MKI67* in all samples.

The low expression of *ALB* (a marker for mature hepatocytes) and the high expression of *TTR* and *GPC3* (markers for hepatocyte progenitors) combined with the high expression of *HNF1B* indicate that the bulk of the cells in the HP samples are hepatocyte progenitors rather than mature hepatocytes or endoderm cells, although there are likely some endoderm cells and immature hepatocytes in the sample (Hay et al., 2008; Mallanna & Duncan, 2013). Similarly, the combined expression of *PDX1* and *NKX6-1* and the low expression of *NEUROG3* (a marker of endocrine progenitors which differentiate from pancreatic progenitors) in the PP samples indicates that these primarily contain pancreatic progenitors but likely contain some endocrine progenitors and endoderm cells (Cogger et al., 2017; Korytnikov & Nostro, 2016).

Notably, HP and PP are closely related cell types that are derived from the same lineage. Indeed, heterogeneous multipotent progenitors can contribute to both the adult liver and adult pancreas in mice (Willnow et al., 2021). Progenitors that express *PDX1* (often used as a marker for the pancreatic lineage) can differentiate into hepatocytes (Willnow et al., 2021). As a result, some overlap in the transcriptomic signature of both cell types is expected and we cannot rule out that the HP samples contain cells that could differentiate into pancreatic cells or that the PP samples contain cells that could differentiate into hepatocytes. However, the expression of *NKX6-1* and *GP2*, markers for pancreatic progenitors, in the PP samples but not the HP samples indicates that these two populations of cells are distinct. Overall, the similarity of PP and HP likely explains the lower number of cell type-specific genes and genes showing cell type-specific ASE for these cell types. This similarity does not alter the conclusions presented in the main text.

Agoglia, R. M., Sun, D., Birey, F., Yoon, S.-J., Miura, Y., Sabatini, K., Pașca, S. P., & Fraser, H. B. (2021). Primate cell fusion disentangles gene regulatory divergence in neurodevelopment. *Nature*, *592*(7854), 421–427. htps://doi.org/10.1038/s41586-021-03343-3

Burridge, P. W., Matsa, E., Shukla, P., Lin, Z. C., Churko, J. M., Ebert, A. D., Lan, F., Diecke, S., Huber, B., Mordwinkin, N. M., Plews, J. R., Abilez, O. J., Cui, B., Gold, J. D., & Wu, J. C. (2014). Chemically defined generation of human cardiomyocytes. *Nature Methods*, *11*(8), 855–860. htps://doi.org/10.1038/nmeth.2999

Chal, J., Al Tanoury, Z., Hestin, M., Gobert, B., Aivio, S., Hick, A., Cherrier, T., Nesmith, A. P., Parker, K. K., & Pourquié, O. (2016). Generation of human muscle fibers and satellite-like cells from human pluripotent stem cells in vitro. *Nature Protocols*, *11*(10), 1833–1850. htps://doi.org/10.1038/nprot.2016.110

Cogger, K. F., Sinha, A., Sarangi, F., McGaugh, E. C., Saunders, D., Dorrell, C., Mejia-Guerrero, S., Aghazadeh, Y., Rourke, J. L., Screaton, R. A., Grompe, M., Streeter, P. R., Powers, A. C., Brissova, M., Kislinger, T., & Nostro, M. C. (2017). Glycoprotein 2 is a specific cell surface marker of human pancreatic progenitors. *Nature Communications*, *8*(1), 331. htps://doi.org/10.1038/s41467-017-00561-0

Hay, D. C., Zhao, D., Fletcher, J., Hewit, Z. A., McLean, D., Urruticoechea-Uriguen, A., Black, J. R., Elcombe, C., Ross, J. A., Wolf, R., & Cui, W. (2008). Efficient Differentiation of Hepatocytes from Human Embryonic Stem Cells Exhibiting Markers Recapitulating Liver Development In Vivo. *Stem Cells*, *26*(4), 894–902. htps://doi.org/10.1634/stemcells.2007-0718

Korytnikov, R., & Nostro, M. C. (2016). Generation of polyhormonal and multipotent pancreatic progenitor lineages from human pluripotent stem cells. *Methods*, *101*, 56–64. htps://doi.org/10.1016/j.ymeth.2015.10.017

Mallanna, S. K., & Duncan, S. A. (2013). Differentiation of hepatocytes from pluripotent stem cells. *Current Protocols in Stem Cell Biology*, *26*, 1G.4.1-1G.4.13. htps://doi.org/10.1002/9780470151808.sc01g04s26

Maury, Y., Côme, J., Piskorowski, R. A., Salah-Mohellibi, N., Chevaleyre, V., Peschanski, M., Martinati , C., & Nedelec, S. (2015). Combinatorial analysis of developmental cues efficiently converts human pluripotent stem cells into multiple neuronal subtypes. *Nature Biotechnology*, *33*(1), 89–96. htps://doi.org/10.1038/nbt.3049

Mickelsen, L. E., Bolisety, M., Chimileski, B. R., Fujita, A., Beltrami, E. J., Costanzo, J. T., Naparstek, J. R., Robson, P., & Jackson, A. C. (2019). Single-cell transcriptomic analysis of the lateral hypothalamic area reveals molecularly distinct populations of inhibitory and excitatory neurons. *Nature Neuroscience*, *22*(4), 642–656. htps://doi.org/10.1038/s41593-019-0349-8

Sharma, R., Khristov, V., Rising, A., Jha, B. S., Dejene, R., Hotaling, N., Li, Y., Stoddard, J., Stankewicz, C., Wan, Q., Zhang, C., Campos, M. M., Miyagishima, K. J., McGaughey, D., Villasmil, R., Matapallil, M., Stanzel, B., Qian, H., Wong, W., … Bharti, K. (2019). Clinical-grade stem cell-derived retinal pigment epithelium patch rescues retinal degeneration in rodents and pigs. *Science Translational Medicine*, *11*(475), eaat5580. htps://doi.org/10.1126/scitranslmed.aat5580

Willnow, D., Benary, U., Margineanu, A., Vignola, M. L., Konrath, F., Pongrac, I. M., Karimaddini, Z., Vigilante, A., Wolf, J., & Spagnoli, F. M. (2021). Quantitative lineage analysis identifies a hepato-pancreato-biliary progenitor niche. *Nature*, *597*(7874), 87–91. https://doi.org/10.1038/ s41586- 021-03844-1
